# Decoding acute neuroinflammatory states from the 3D architecture of *in vitro* microglia

**DOI:** 10.64898/2025.12.19.695376

**Authors:** Anthoula Chatzimpinou, Johanna Huber, Xenia Podlipensky, James-Lucas Cairns, Ayse Erozan, Jian-Hua Chen, Valentina Loconte, Mark A Le Gros, Carsten Hopf, Carolyn Larabell, Venera Weinhardt

**Author notes:** Correspondence author contact details, Postal address: Institute of Microstructure Technology IMT, Hermann-von-Helmholtz-Platz 1, 76344, Eggenstein-Leopoldshafen, Germany.

## Abstract

Microglia, core brain immune defenders, rapidly polarize into pro- or anti-inflammatory states that shape neuronal survival during acute brain inflammation. Yet, how these inflammatory states are encoded at the native cellular level remains unclear. While microglial states associate with specific molecular and organelle markers, it is unknown whether their cellular architecture integrates robust metabolic and structural features. Here, we quantitatively decode inflammatory states from the 3D architecture of individual *in vitro* mouse microglia (BV-2). Using soft X-ray tomography on established BV-2 polarization, we identify coordinated intracellular organization distinguishing homeostatic, pro-inflammatory, and anti-inflammatory cells, including characteristic lipid droplet–endosome architectures. We further link lipid-endosome reorganization with mTORC1 pathway, by functionally implicating sestrin-2 in promoting anti-inflammation. Finally, we resolve the time-dependent remodeling of lipid droplet 3D profiles by their distinct lipidomic composition across inflammatory states. Overall, our work enables deciphering microglia states in disease-relevant models, with potential to ultimately understand brain immunity.

## Introduction

Neuroinflammation is a central histopathological hallmark of neurodegenerative diseases, manifesting from acute to chronic clinical cases^1^. Microglia, the brain’s resident macrophages, rapidly adopt distinct functional states during the primary immune response, acting as the first line of defense against toxic stimuli^2^. Their cytotoxic activity contributes to neurodegenerative diseases such as ischemic stroke and Alzheimer’s, with histopathological evidence highlighting the role of microglia-driven neuroinflammation in disease progression and treatment outcomes^3–5^.

Considering of the brain’s intricate architecture^6,7^, microglia display significant regional heterogeneity in both morphology and molecular signatures^8–10^. This phenotypic diversity poses challenges in consistently characterizing microglial populations, especially as they shift between functional states^8,9^. A widely adopted, though simplified, framework categorizes microglia into three states: a homeostatic baseline (M0), a pro-inflammatory (M1), and an anti-inflammatory (M2) phenotype^11^.

The M2 phenotype is activated by specific biochemical signals that induce different microglial subtypes, classified as M2a, M2b, M2c, and M2d^12^. Among those, M2a is the most prevalent and is stimulated by interleukin-4 (IL-4) and interleukin-13 (IL-13), leading to the release of arginase-1 (ARG-1), which promotes anti-inflammatory properties of microglia^13^. This terminology reflects responses to different stimuli, with M1 microglia promoting the release of inflammatory cytokines, like inducible nitric oxide synthase (iNOS), and contributing to neuronal damage, while M2 microglia act as neuroprotectants and resolve neuroinflammation^11,14^. Although these functional states have been explored across various levels, from molecular profiles to cellular morphology^8^, an integrative approach that captures multilevel transitions within cells has yet to be fully established.

Recent advances in 3D imaging have introduced the concept of “whole-cell architecture”^15^. This approach provides detailed structural insights into cellular ultrastructure, encompassing all organelles larger than 100 nm, such as the nucleus, mitochondria, and lipid droplets^16^, across cell types and states^17^. By providing information on the functional interconnectivity of all organelles within cells, such approaches enable us to understand cell biology under pathological conditions at an unprecedented level.

Among the 3D imaging techniques capable of revealing whole-cell architecture, soft X-ray tomography (SXT) stands out as a biochemically sensitive and quantitative imaging modality^15,16,18^. During tomographic acquisition, individual cells are exposed at all angles under X-rays in fully hydrated, cryogenically preserved conditions, at a spatial resolution of 25-60 nm^19^. Leveraging the "water window", a spectral range where water is relatively transparent but carbon- and nitrogen-containing biomolecules strongly absorb X-rays^20^, SXT generates natural contrast without the need for staining or labeling^21^. This intrinsic contrast allows voxel-based quantification of the linear absorption coefficient (LAC, μm⁻¹), reflecting the local biochemical content within different organelles^18,22,23^. By capturing subtle biochemical variations at high spatial resolution, SXT enables the investigation of coordinated structural, metabolic, and molecular processes in transitioning cells in numerous scientific cases^16^. Different LAC-based analytical approaches have provided diverse readouts, including tracking nucleotide base conversions^16^, linking transcriptional signatures to developmental stages in olfactory neurons^24^, modeling and classifying insulin granules in pancreatic β cells under varying glucose conditions^25,26^, and charting the cell cycle of the recently discovered nitroplast in symbiotic algae^27^.

In our work, we combine SXT imaging with molecular and biochemical characterization to examine 3D cell architecture at different neuroinflammatory states of individual microglia, using an *in vitro* polarization model - mouse microglia cell line (BV-2)^28^. First, we molecularly confirm the polarization robustness of BV-2 cells by activating them with state-specific triggers, like lipopolysaccharides (LPS) for M1 and IL-4 for M2a state. Based on this established *in vitro* model, we probe with SXT imaging 3D structural profiles of individual microglia during the transition from homeostatic (M0) to pro-inflammatory (M1) or anti-inflammatory type-a (M2a) state. To further validate the state-related structural signatures over microglia polarization, we integrate the 3D structure of cells obtained with SXT imaging with other approaches, like correlative fluorescence and lipid mass spectrometry, targeting potential intracellular processes that further help to understand microglia-driven neuroinflammation.

## Results

### BV-2 cells are robustly polarized to the M1 or M2a state at the single-cell level

To investigate the cell architecture of polarized microglia, we first established a robust and efficient *in vitro* model of M1/M2a activation using the immortalized murine microglial cell line, BV-2^29^. This cell line is the most widely employed *in vitro* model, shown to replicate key genetic (e.g., interferon-gamma (IFN-γ) and iNOS expression) and morphological traits (e.g., cell body ramification) of primary microglia when activated during neuroinflammation^28,30^. While BV-2 cells exhibit lower transcription levels of transforming growth factor *Tgf-*β and reduced expression of microglial homeostatic markers, such as the purinergic receptor *P2ry12*, they demonstrate robust responses to immune stimuli, including lipopolysaccharides (LPS) and interleukin-4 (IL-4)^31–34^. Overall, previous work suggests that BV-2 cells are a reliable model for studying acute neuroinflammation^28,30,35^.

To establish robust polarization of microglia cells to M1 or M2a state, we have assessed the polarization rate over the cell population treated with varying concentrations of LPS (0.1–10 μg/mL) or IL-4 (20–40 ng/mL) over 6 and 24 hours, based on iNOS and ARG1 expression levels. The BV-2 cells were cultured for two weeks to maintain their homeostatic (M0) state (Figure 1a). Upon reaching 70-80% confluency, pro-inflammatory (M1) or anti-inflammatory (M2a) states were induced using LPS or IL-4 and evaluated by bulk protein expression using western blot at different time points (Figure 1b-c). We have chosen an intermediate point of analysis at 6 hours for both LPS and IL-4 as the inflammatory-related proteins become more stable after 24 hours post-inflammation, while the first 6-8 hours are early enough to capture initial microglia responses^14,36^. From western blot images (Figure 1b-c; Figure SI1), it is evident that both M1 and M2a polarization states depend on the concentration of the respective drugs; however, the expression of ARG1 is enriched only at 24 hours. This suggests different kinetics in the activation of M1 and M2a states (Figure 1b-c), consistent with previous studies^37,38^.

**Figure 1.**
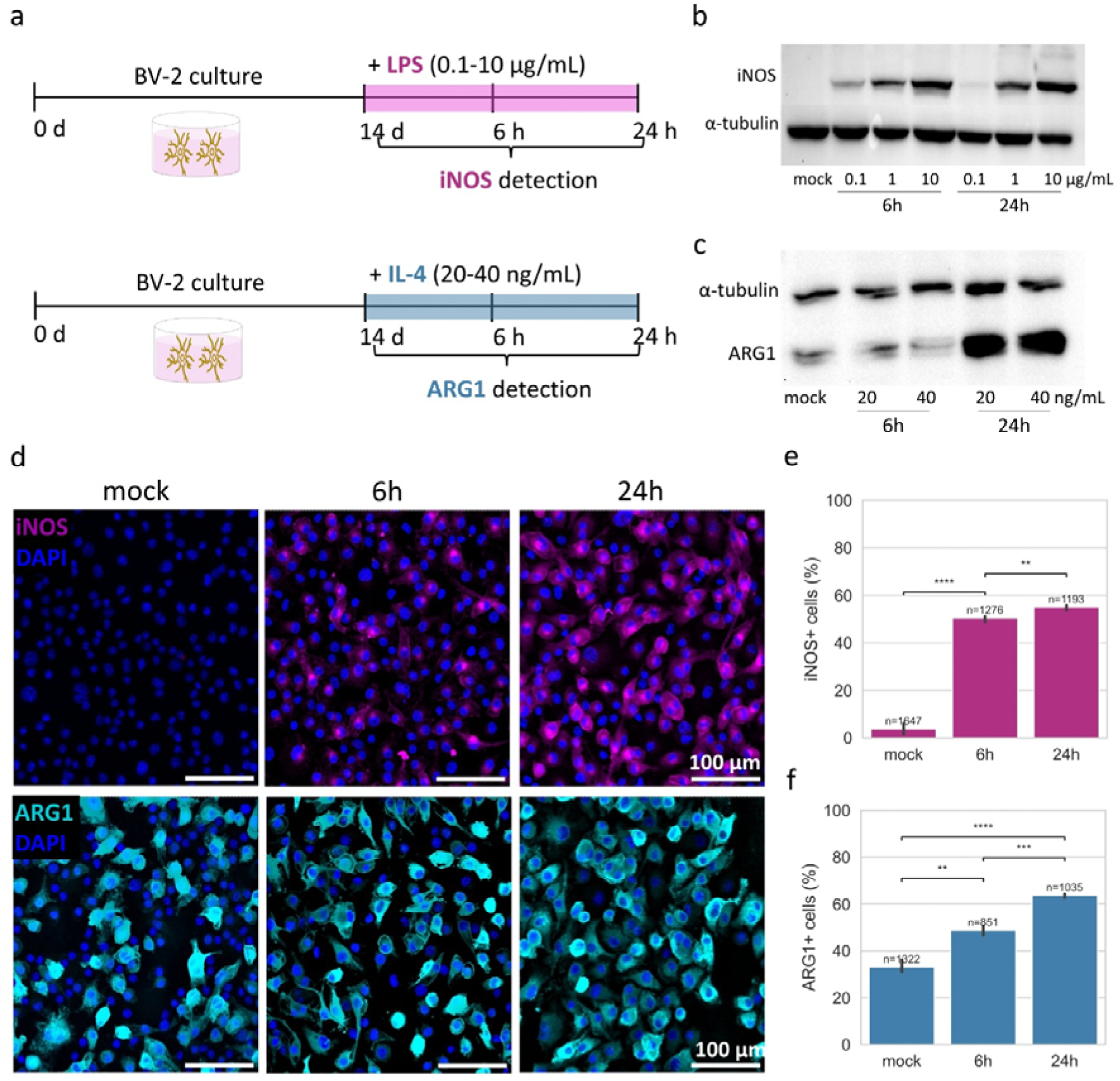
Robust polarization of BV-2 cells to M1 and M2a state. **(a)** Schematic of *in vitro* setup for M1 or M2a activation with LPS or IL-4 at 6 and 24h. Activation rates were analyzed with Western blot for **(b)** iNOS and **(c)** ARG1 expression, using α-tubulin as a control. **(d)** Immunofluorescence images of BV-2 cells for iNOS and ARG1 in mock and LPS/IL-4 treated conditions. Confocal images were acquired with a 20x objective; scale bar = 100 μm. Quantified proportions of **(e)** iNOS^+^ and **(f)** ARG1^+^ cells were based on cytoplasmic intensity in 3 images per condition. Statistical analysis: t-test, p-values: ns (0.05 < p ≤ 1.00), * (0.01 < p ≤ 0.05), ** (0.001 < p ≤ 0.01), *** (0.0001 < p ≤ 0.001), **** (p ≤ 0.0001). Sample size, i.e. number of analyzed cells, (n): mock (no LPS)=1647, 6h (LPS)=1276, 24h (LPS)=1193, mock (no IL-4)=1322, 6h (IL-4)=851, 24h (IL-4)=1035.

Considering that for optimal subcellular analysis with SXT, most cells exhibit structural changes, we selected the highest concentrations of LPS (10 μg/mL) and IL-4 (40 ng/mL) that maximize iNOS and ARG1 expression over 24 hours. To analyze individual cells, we first confirmed robust single-cell activation in cell culture. Immunofluorescence imaging of iNOS and ARG1, with co-staining of the nuclear and plasma membrane, was used to identify and automatically segment expression levels in individual cells (Figure SI2). This enabled us to quantify M1/M2a polarization at the population level at 6 and 24 hours. In the LPS-treated culture, 50% (n=1276, p=6.260 × 10⁻□) of cells were iNOS^+^ at 6 hours. As iNOS expression increased, the proportion of iNOS^+^ cells slightly rose (n=1193, p=9.070 × 10⁻³) by 24 hours (Figure 1d-e). Since ARG1 is ubiquitously expressed under mock conditions, 25% of cells were ARG1-positive before the treatment. The addition of IL-4 steadily increased the proportion of ARG1^+^ cells, reaching 65% (n=1035, p=5.674 × 10⁻□) by 24 hours (Figure 1d-f).

Altogether, we achieved robust activation of BV-2 cells with high concentrations of IL-4 (40 ng/mL) and LPS (10 μg/mL), with most cells (more than 60%) polarized into an M1 or M2a state, providing sufficient cells for probing single-cell structure by soft X-ray tomography.

### SXT analysis detects intracellular correlation upon microglia activation

Soft X-ray tomography (SXT) data collection is rich in structural information on individual organelles, their spatial interactions, and their chemical states in different pathological conditions^18,25,39^. Most approaches use the classical comparison of two features at the same time, or a comparison of gross morphological parameters^18,21^. Recently, SXT has been used for the clustering of individual cells based on particular organelle structural alterations, like insulin granules of pancreatic cells, in different extracellular conditions^22,26^. Given the potential difference in activation states of individual cells and the complexity of microglia polarization, we have chosen to develop a quantitative SXT analysis that would reflect the organization and chemical state of organelles within entire cells at the cell population level^15^.

For SXT imaging, we followed an established procedure for sample preparation^19^. The BV-2 cells in M0, M1 and M2a states at 6 and 24 hours were imaged in 3D with 50nm spatial resolution, yielding 3D ultrastructural data of single cells (Figure 2a)^19,40^. Each field of view (FOV) of the SXT depicts the entire cell volume, including visible ultrastructure, as illustrated by a 2D virtual slice from a tomogram (Figure 2b). Due to the increased BV-2 cell size in most activation states, three to four tomograms were combined to ensure the entire cell volume was included (Figure SI3). Before probing individual organelles, the whole cell volume was segmented using the ACSeg model^41^, enabling the separation of cells from the background and specimen holder (Figure SI3). Based on specific LAC values and known morphology, the different organelles, such as the nucleus, mitochondria, lipid droplets, and endosomes, were manually segmented. For quantitative analysis, we selected 3D parameters of individual organelles, including volume (V, μm³), surface area (A, μm²), surface area-to-volume ratio (A/V), and biochemical density (LAC, μm ¹) (Figure 2b-c, Figure SI3). Both their absolute and relative (to the whole cell) values were used for further analysis. In Figure 2b and 2d, exemplary results of such segmentation for M0, M1, and M2a states are shown at different activation times. Such 3D renderings of segmented organelles illustrate their gross morphology and spatial distribution within the cell, thereby building a visual atlas of whole cells (Figure 2b).

**Figure 2.**
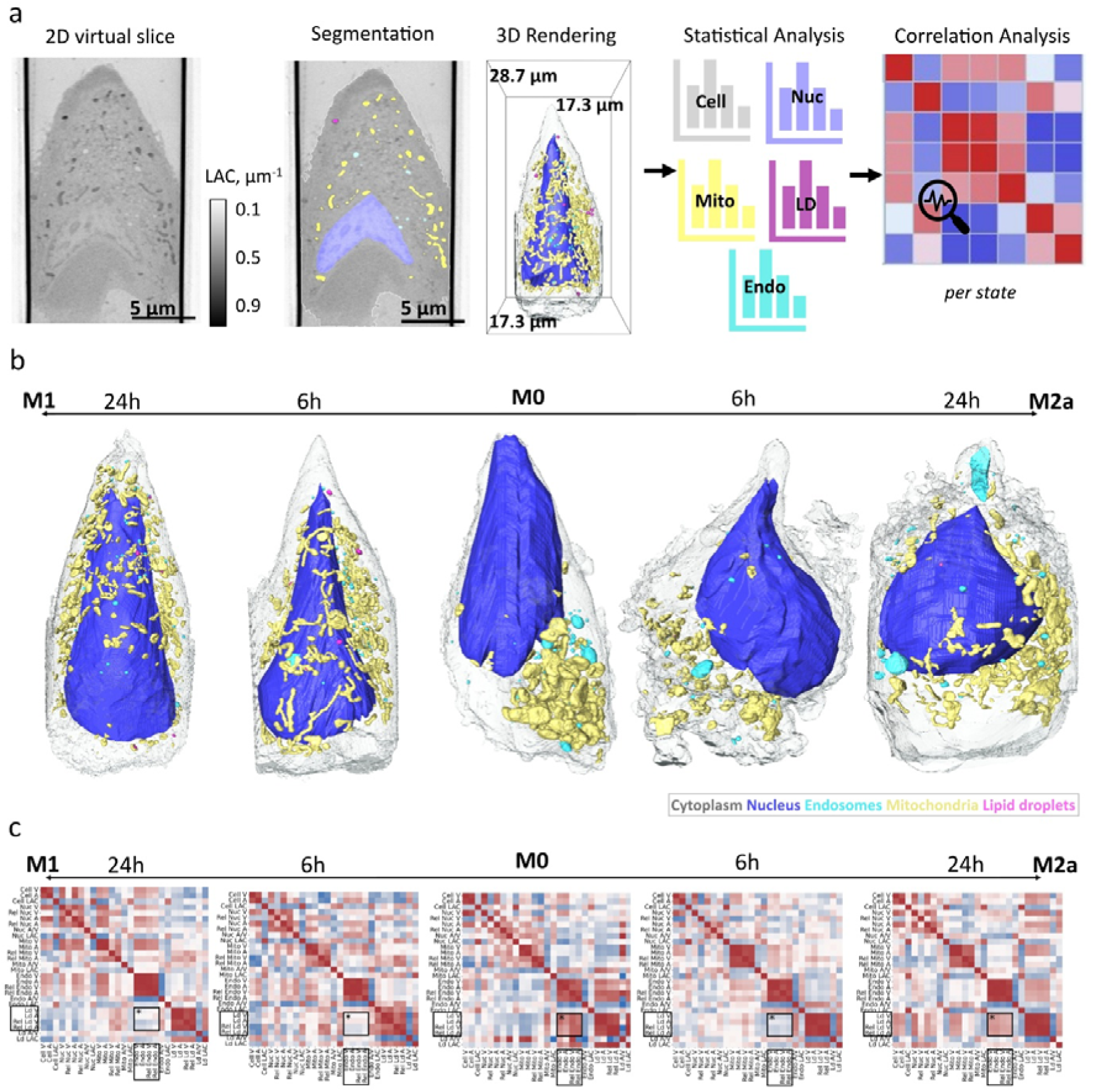
SXT analysis detects intracellular correlation upon microglia activation. **(a)** 3D analysis is based on the segmentation of different organelles of individual cells, assigning to specific intensity values (LAC, colourmap set at 0.1-0.9 μm^-1^). Here: whole cell in white outline, nucleus in blue, mitochondria in yellow, endosomes in cyan, and lipid droplets in magenta. The scale bar of the 2D virtual slice is 5μm. 3D rendering of segmented organelles shown in the bounding box, showing the physical size of the tomogram (x:17.3 μm, y:28.7 μm, and z:17.3 μm). Structural analysis describes inter-organelle patterns based on Pearson’s correlation analysis of 3D morphological features of individual cells per state per time point. **(b)** 3D renderings of individual cells representing microglia at each state (M1/M2a or mock) at 6 and 24 hours. Lipid droplet and endosome renderings are dilated by a factor of 2.0 for better visualization. **(c)** Cross-correlative matrices show the organelle co-rearrangements over microglia M1/M2a polarization. Pearson’s coefficient (R); -0.75 (blue) to +1 (red). Sample size (n); 13 cells (M0), 13 cells (M1 6h), 13 cells (M1 24h), 12 cells (M2a 6h), and 13 cells (M2a 24h). Black boxes with an asterisk (*) indicate the LD-Endo volumetric changes for each cell condition.

To gain a comprehensive understanding of the ongoing anatomical changes that microglia undergo during their polarization to the M1 or M2a state, we have developed an evaluation pipeline that probes the orchestration of organelles in pair-wise mode in color-coded matrices using Pearson’s correlation method (Figure 2c). The pipeline begins by evaluating the relationship between organelle gross morphological changes, based on their mean absolute and relative 3D parameters in individual microglia cells when polarized to an M1 or M2a state over time (Figure 2a,c). The colormap helps to visualize the linear relationship between two parameters, where red represents Pearson’s correlation coefficient (R) values closer to +1, indicating a positive relationship, and blue represents values closer to -0.75, pointing to a negative relationship (Figure 2c). Light colors on the red-blue spectrum, closer to white, show no linear relationship between the parameters (Figure 2c).

The developed pipeline should provide insights into the morphological relationships between organelles during microglial polarization. We, therefore, applied this pipeline to microglia at different activation states, focusing on the organelle relationships that show differences upon polarization to an M1 or M2a state compared to the M0 state. Apart from the expected changes, such as the nucleus (Nuc)-to-cell volume ratio, which increases over pro-inflammatory activation due to cell body expansion^9^, intracellular patterns involving lipid droplets (LD) and endosomes (Endo) were the most prominent (Figure 2c).

Increased LD accumulation coincides with thinner mitochondria in M1, whereas lipid droplets and mitochondria expand their respective volumes in M2a, indicating distinct metabolic adaptations (Figure 2b,c). Mitochondrial fission has been previously correlated with their dysfunctional respiration and the initiation of metabolic processes, such as the TCA cycle^42,43^. Concurrent studies link this phenotype with the excess of lipid droplets in M1-inflammatory microglia through molecular interactions in the LD-mitochondria axis^44^. Similarly, the dynamics of LDs and mitochondria also vary across states in our study (Figure 2c). Mitochondria show a fused phenotype with larger volumes in the M0 and M2a states, compared to a thinner, fissured phenotype in M1, which has been previously correlated with pro-inflammatory microglia^43,45^. In the M0 state, lipid droplets and endosomes are positively correlated, a relationship that diminishes at early activation (6h) in both M1 and M2a states. By 24 hours, this correlation becomes moderately negative in M1 but remains strongly positive in M2a, suggesting a temporary blockage in endosome maturation in the M1 state due to the presence of lipid droplets.

Based on the whole-cell analysis of phenotypic changes in single microglia upon activation by SXT, we investigated the regulation of the endosomal lipid pathway in more detail.

### LD-to-Endosome association indicates M1/M2a-specific metabolism

Leveraging the outcome of our structural analysis, we focused on metabolism-related co-altering patterns, starting with the lipid droplet (LD)-endosome (Endo) relationship. As previously discussed, LD presence over activation leads to distinct morphological changes in other organelles, such as mitochondria or endosomes (Figure 2d-e). Mitochondria have consistent and distinct LAC (0.27-0.33 μm^-1^ in BV-2) from other organelles inside the cells and have been widely analyzed for their structure in previous SXT-based studies^21,39,46^. In the current analysis, we also describe endosome 3D morphology, due to the phagocytic properties of microglia cells^47^. Depending on their maturation stage towards lysosomes, endosomes share diverse phenotypes inside the cells, and their characterization relies on their molecular expression and overall structure^48^. Based on previous nanostructure analyses of endosomes from electron microscopy^49,50^, we segment endosomes of different sizes and shapes (round or irregular, small or big, with or without a rim) and measure their LAC, which is close to the extracellular aqueous space, at 0.16-0.19 μm^-1^ (Figure 3a). In contrast to endosomes, lipid droplets are overall round, smaller, and biochemically denser, with LACs ranging between 0.35 and 0.7 μm^-1^ (Figure 3a).

**Figure 3.**
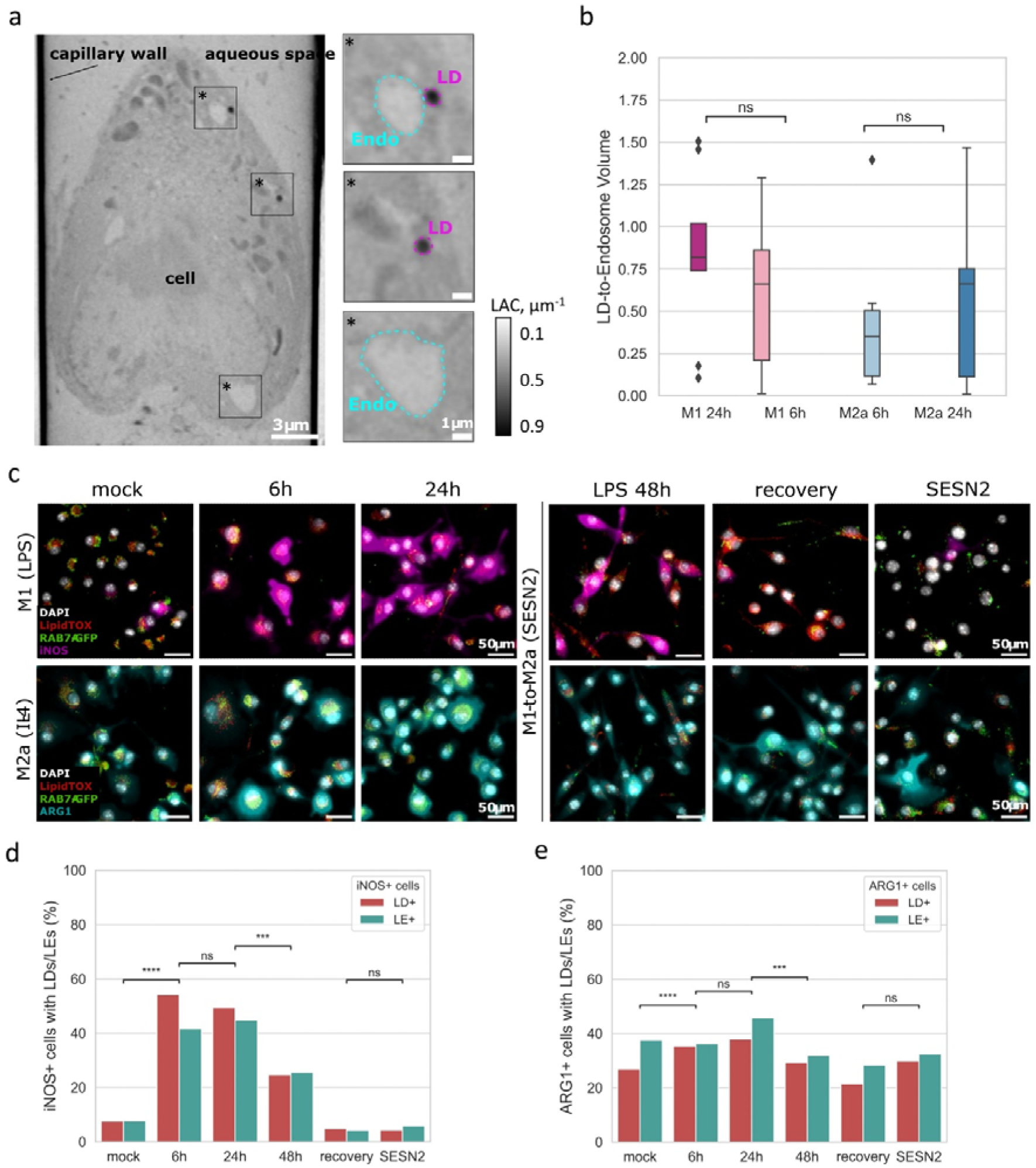
LD-to-Endo pattern indicates M1/M2a polarization-associated metabolism. **(a)** 2D virtual slice of an SXT-imaged cell with all the intracellular information shown. The scale bar is 3 μm. Three boxes with asterisk (*) zoom in on the endosomes (cyan outline) and lipid droplets (magenta outline), sharing different morphology and LAC values. The scale bar is 1 μm, and the LAC range is set at 0.1-0.9 μm^-1^. **(b)** Relative LD-to-endosome volume is slightly bigger in M1 24h than in the rest of the states in the SXT datasets. Statistical analysis performed in t-test independent samples, p-value (p) ns: 0.05 < p ≤ 1.00, (*): 0.01 < p ≤ 0.05, (**): 0.001 < p ≤ 0.01, (***): 0.0001 < p ≤ 0.001, (****): p ≤ 0.0001. The sample size (n) is: 13 cells (M1 6h), 13 cells (M1 24h), 12 cells (M2a 6h), and 13 cells (M2a 24h). **(c)** FOVs from light microscopy images show the presence of organelle markers (Les-green, LDs-red) in the immunofluorescent for iNOS (magenta) and ARG1 (blue) cells. Images were sequentially acquired as z-stacks at the same FOVs before and after immunofluorescence, using an air-immersion 20x objective on a wide-field fluorescent microscope (ACQUIFER machine). Scale bars are set at 50 μm. **(d-e)** Quantification shows the relative presence of LDs and Les in iNOS^+^ and ARG1^+^ cells over LPS/IL-4 treatment and LPS recovery with or without SESN2 (12.5 ng/mL). Statistical analysis performed in t-test independent samples, p-value (p) ns: 0.05 < p ≤ 1.00, (*): 0.01 < p ≤ 0.05, (**): 0.001 < p ≤ 0.01, (***): 0.0001 < p ≤ 0.001, (****): p ≤ 0.0001. For iNOS analysis, the sample size (n) is mock n=475, 6h (LPS) n=231, 24h (LPS) n=272, 48h (LPS) n=379, 48h (recovery) n=475 and 48h (recovery + SESN2 12.5 ng/mL) n=266 in 3 FOVs per condition. For ARG1 analysis, mock n=390, 6h (IL-4) n=461 24h (IL-4) n=496 cells, 48h (LPS) n=379, 48h (recovery) n=475 and 48h (recovery + SESN2 12.5 ng/mL) n=266 in 3 FOVs per condition.

From the 3D representation of mock and polarized microglia (Figure 2b), there is a difference in size over activation time for both types of polarization; in M1, smaller and rounder endosomes are frequently found, whereas in M2a, bigger and more mature endosomes with a rim, possibly fused with lysosomes. To pair this phenotypic change with the presence of LDs over polarization, we assessed the volumetric ratio balance between the two organelles (Figure 3b). While a 1.0 LD-Endo ratio is preserved during activation to the M2a state, in M1, there is a clear LD volumetric dominance over the endosomes (Figure 3b). We suggest that this effect of LDs on endosomes relates to a blockage of endosome maturation occurring during M1 polarization.

Concentrating on possible mechanistic relationships associated with endosome maturation and lipid droplets that signify M1 or M2a polarization, we triggered the M1-to-M2a transition using sestrin 2, which has been previously shown to block mTORC1 signaling and promote lysosome maturation in BV-2 cells under ischemic stroke conditions^51^. For molecular verification of activation state related to LDs or endosomes accumulation, we developed a correlative fluorescence pipeline. To assess LD-late endosome (LE) changes at mock, M1/M2a, and M1-to-M2a transition stages over time, we first imaged BV-2 cells labelled for LD (LipidTOX) and LE (Rab7A-GFP), followed by immunostaining and imaging of iNOS (M1) and ARG1 (M2a) levels of the same cells (Figure 3c, Figure SI4).

Comparative analysis of LD- and LE-containing iNOS^+^ and ARG1^+^ cells reveals a dynamic shift in microglia metabolism during polarization (Figure 3d-e). In early M1 activation, LDs dominate, with a gradual change towards LE-containing iNOS^+^ cells, though LDs remain prevalent at 24h (Figure 3d). This trend resembles the LD-Endo volumetric ratios obtained by SXT (Figure 2b), supporting the transient LD presence in M1 cells. The M2a-polarized ARG1^+^ cells on the other side show a lower LD-to-LE ratio, which is consistent across both IL-4 treatment and mock conditions (Figure 3e). During the M1-to-M2a transition with SESN2-assisted recovery, LE-containing ARG1^+^ cells dominate the M2a population, maintaining the mock cell LD-LE ratio (Figure 3e). Yet, SESN2 treatment post-LPS challenge enhances LE abundance in both iNOS^+^ and ARG1^+^ cells, suggesting a metabolic shift favoring LD breakdown and endosomal maturation. However, the abundance of LE is similar to recovery from LPS by exchanging cell media with no significant change towards M2a polarization (Figure SI5, panels a-c).

In summary, we suggest that the LD-LE pattern aligns with LD-endosome correlations in SXT imaging, reinforcing the distinct LD-LE shift in M1 and M2a states. Although SESN2 enhanced the M2a-related LD-LE balances post M1 polarization in the overall BV-2 population, in our study, it did not effectively promote the molecular shift towards ARG1 expression.

### M1/M2a-polarized microglia exhibit distinct LD structure and lipid profiles

Accumulative lipid droplets (LDs) are associated with pro-inflammatory microglia, making them potential inflammation markers^52^. Therefore, LD profiling is currently being intensively examined in neurodegenerative tissues to connect the LD compositional difference to inflammation^53,54^. In our SXT findings, LDs are present in 20% of M0 microglia at minimal volumes, and their presence and size increase in the majority of M1 and M2a cells at 6 and 24 hours (Figure SI5, panel d), consistent with the previously established relationship between LD accumulation and inflammation^55^. Since LD volume remains unchanged across inflammatory states (Figure SI5, panel e), we exclude M0 from further LD analysis.

Apart from structural information, SXT data provides insights into chemical composition, which has been used to detect the maturation stages in insulin granules^25^, herpes viral maturation^23^, and oxidative state of mitochondria upon infection^39^– all via LAC shifts. Consequently, we assessed the biochemical content of individual LDs based on their mean LAC (0.35–0.7 μm⁻¹) at the single microglia over M1/M2a polarization (Figure 4a). The 3D renderings show that LDs exhibit higher X-ray absorption in late M1 activation and lower, more homogeneous absorption in M2a (Figure 4a). The LAC analysis in correlation to LD volume shows that LD phenotypes derived from M1- and M2a-polarized cells diverge at 24 hours but overlap at 6 hours, suggesting distinct lipid compositions between pro- and anti-inflammatory activation over time (Figure 4b).

**Figure 4.**
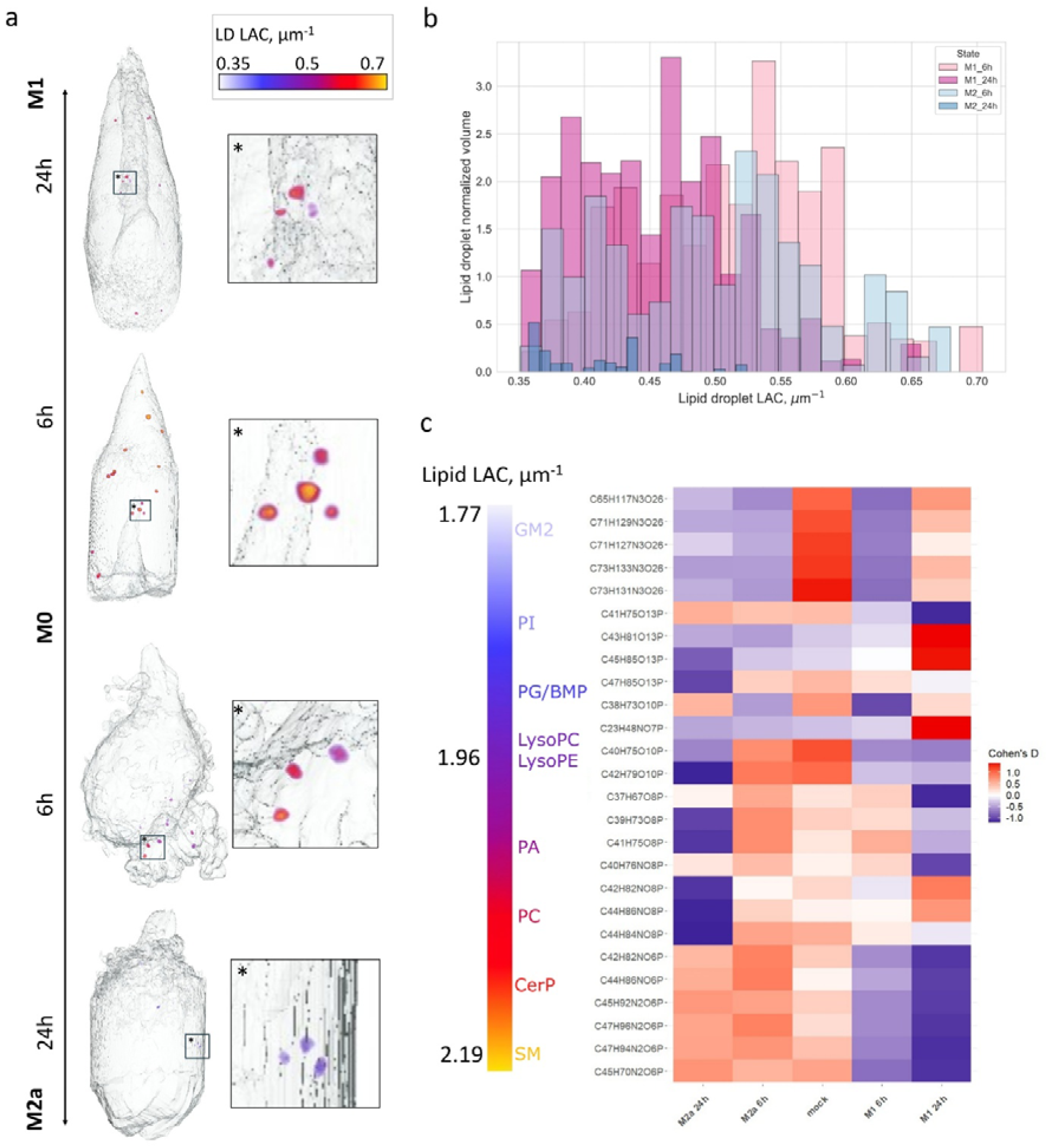
M1/M2a-polarized microglia display distinct LD 3D structure and lipid composition. **(a)** 3D renderings of representative M1 and M2a polarized BV-2 cells display the mean linear absorption coefficient (LAC) of individual lipid droplets (LDs), visualized using a colormap ranging from 0.35 to 0.7 μm⁻¹ (white/blue to yellow). **(b)** LDs demonstrate distinct 3D morphological distributions characterized by normalized volume and LAC across the M1/M2a polarization spectrum, with M2a states shown in light blue/blue (6 h and 24 h) and M1 states in light pink/pink (6 h and 24 h). **(c)** MALDI-mass spectrometry analysis reveals the relative abundance of 26 individual lipids (characterized on MS2 level) across mock, M1, and M2a polarized BV-2 cells at 6 and 24 hours. Lipids are sorted from top to bottom according to their estimated LAC (μm⁻¹) at 520 eV, derived from their known chemical formulas, and are color-coded by lipid subclass. The colormap ranges from 1.77 to 2.19 μm⁻¹ (white/blue to yellow). All lipids plotted here, show significant differences compared to mock after at least one of the treatments, which was assessed by independent t-tests (p-value <0.05). Cohen’s d effect size shows intensity differences, where strong positive (red) indicates d ≥ 1.0, neutral (white) d ≈ 0.0, and strong negative (blue) d ≤ -1.5, based on three independent replicates of 1 million cells per condition.

To determine whether differences in LD structure are associated with modulation of overall lipid composition among the M1/M2a polarized microglia, we performed MALDI mass spectrometry^56^ on the bulk BV-2 population under control and LPS/IL-4-treated conditions at 6 and 24 hours. Based on our lipidomic analysis of each condition, we identified 26 individual lipid species, categorized into 8 lipid subclass groups (Table 1 and Figure 4c). These groups contain one or two lipid subclasses, which were all validated by tandem mass spectrometry (MS^2^ level annotation, Table SI1 and Figures SI6-13). To correlate the lipid abundance in the cell population with the LAC measured in individual cells (Figure 4a), we calculated and sorted the LAC of the identified lipid species, using their known chemical sum formulas at a fixed X-ray energy of 520 eV^57^ (Table 1 and Figure 4c). In this analysis, the lipid subclass groups exhibited distinct X-ray absorption ranges, which could be leveraged for their classification (Table 1). Among sphingolipids, gangliosides (GM2s) and sphingomyelins (SMs) exhibited the lowest (1.77 μm^-1^) and highest (2.19 μm^-1^) LACs, respectively, while diverse phospholipid groups absorbed X-rays across a broad intermediate range (1.82–1.99 μm⁻¹). Regarding LD-related species, our whole-cell MALDI MS lipidomic analysis did not detect any neutral lipids, such as triacylglycerols (TAGs) or cholesteryl esters (CEs), commonly stored in LDs^58^. This likely reflects their relatively small contribution to the total cellular mass in BV-2 cells, as previously shown by volumetric analyses with SXT (Figure 3b & Figure SI 5e), which might limit their detection by MALDI MS in BV-2 cells, since the current technique has previously identified TAGs in a different microglial cell line treated with LPS^56^.

**Table 1.** Lipid subclasses detected in M1/M2a-polarized and mock BV-2 cells with MALDI-mass spectrometry (Metaspace annotation with false-discovery rate (FDR) <10%) and their associated LAC values, calculated as reported in the Methods (“Cryo soft X-ray tomography”).

| Lipid subclass group (abbreviation) | Lipid class | LAC ($\mu\text{m}^{-1}$) |
| --- | --- | --- |
| Ganglioside (GM2) | Sphingolipid | 1.77-1.82 |
| Phosphatidylinositol (PI) | Phospholipid | 1.82-1.88 |
| Phosphatidylglycerol (PG) or Bis(monoacylglycerol)phosphate (BMP) | Phospholipid | 1.89-1.92 |
| Lysophosphatidylcholine (LysoPC) or Lysophosphatidylethanolamine (LysoPE) | Phospholipid | 1.89 |
| Phosphatidic acid (PA) | Phospholipid | 1.96-1.99 |
| Phosphatidylcholine (PC) | Phospholipid | 2.02-2.04 |
| Ceramide phosphate (CerP) | Sphingolipid | 2.10-2.11 |
| Sphingomyelin (SM) | Sphingolipid | 2.14-2.19 |

Our integrated SXT–MALDI MS analysis revealed distinct lipid profiles characterizing homeostatic (M0) and M1/M2a inflammatory microglia (Figure 4c). Homeostatic M0 cells contain almost all lipid groups, except for phosphatidylinositols (PIs) and lysophosphatidylcholines (LysoPCs) or lysophosphatidylethanolamines (LysoPEs). These classes are uniquely found in M1 cells, with particular LysoPC species showing almost a 2.3-fold increase upon LPS treatment^56^. High enrichment of gangliosides (GM2s) is a defining feature of M0 cells, distinguishing them from reactive M1 or M2a states. While low in GM2, M2a-inflammatory cells share an abundance of sphingolipids similar to M0 cells. Notably, M2a cells shift from diverse enrichment in phospholipid-related groups at 6 hours to a strong enrichment in sphingolipids at 24 hours, reflecting the 3D morphometric changes of lipid droplets under this condition (Figure 4b). In contrast, M1-inflammatory microglia lack highly X-ray-absorbing sphingolipids (GMs or CerPs), although they contain GM2 over 24 hours. At 24 hours, M1 cells exhibit unique phospholipid signatures with divergent LACs, whereas at 6 hours, their lipid diversity is generally low. This suggests transient lipid remodeling under these conditions, which aligns with the 3D architecture profiles shown with SXT (Figure 2c).

Taken together, our findings indicate that LDs undergo distinct 3D morphological shifts between M1 and M2a states over time, associated with lipid remodeling dynamics that characterize the transition from homeostatic to pro- or anti-inflammatory conditions and reflect their whole-cell 3D architecture profiles.

## Discussion

Microglia maintain brain homeostasis and respond to diverse inflammatory stimuli by adopting different activation states^59^. Their activation has been mainly described through the M1/M2 polarization, based on classical markers^11^. While helpful in describing neurodegenerative diseases^11^, this terminology oversimplifies the spectrum of states microglia can adopt in response to context^8^. Here, we introduce 3D cell architecture as a readout that integrates molecular, metabolic, and structural features of two neuroinflammatory states, providing a comprehensive view of *in vitro* microglia polarization and function. By combining soft X-ray tomography (SXT), MALDI mass spectrometry, and molecular assays, we show how organelle and lipid remodeling coordinate during transitions from homeostatic (M0) to pro-inflammatory (M1) or anti-inflammatory (M2a) states.

Anticipating that cryo-SXT imaging at a capillary setup makes direct molecular confirmation challenging, we validated beforehand the dynamics of BV-2 polarization with quantitative immunofluorescence. The temporal dynamics of state-specific marker expression closely resembled those observed in primary microglia, reinforcing the translational relevance of our *in vitro* model^60,61^. The high efficiency of polarization, reaching 55-60% at 24 hours, enabled us to perform single-cell SXT imaging within a heterogeneous cell population. Such heterogeneity is biologically meaningful, reflecting the diversity of microglial responses, even within an *in vitro* controlled experimental set-up.

Regarding data interpretation, for organelles with complex molecular and morphological features, like the endosomes, we developed a downstream validation with fluorescent labeling. To evaluate the endosome maturation over polarization, we targeted only the late endosomes (LEs) by tagging them with the reference Rab7 protein. Since there are more endosome types (e.g., early endosomes, endo-lysosomes, etc) found in microglia and related to their function^62^, we suggest further validation with other, more spatially sensitive techniques, like cryo-electron microscopy.

Our results revealed distinct structural and lipid adaptations from homeostatic to a neuroinflammatory state. In M0 microglia, the minimal presence of LDs compared to late endosomes was accompanied by a high enrichment in GM2 gangliosides, a feature that clearly distinguishes them from M1 or M2a inflammatory states. Gangliosides, which primarily localize within the plasma membrane^63^, have been previously associated with the phagocytic activity of microglia in the ischemic brain^64^. Notably, ganglioside GM1 enhanced the endo-lysosomal maturation and anti-inflammatory responses following pro-inflammatory challenges in BV-2 and primary microglia^65^. More recent findings also highlight the role of genes, like the beta subunit of β-hexosaminidase (*Hexb*), in coordinating the necessary microglia-neuron homeostasis via regulating the ganglioside GM2 in phago-lysosome compartments in microglia^66^. While ganglioside homeostatic functionality and endo-lysosomal positioning are pinpointed by these findings, shifting BV-2 cells to an M2a-inflammatory state negatively affects their presence in those cells, shifting towards other sphingolipid classes, such as sphingomyelins (SMs) and ceramide phosphates (CerPs).

Under anti-inflammatory conditions, BV-2 microglia temporarily accumulate LDs, which are degraded mainly within 24 hours, accompanied by a pronounced enhancement of ARG1 and a balanced lipid droplet–late endosome composition. The M2a cells contained smaller, less dense LDs than M1 and were overall enriched in ceramide phosphates (CerPs) and sphingomyelins (SMs), lipid classes linked to membrane repair, antioxidative, and anti-inflammatory functions^67^. In macrophages, the parallel ceramide biosynthesis to phagosome maturation is mediated by ceramide synthase 2 (Cer2), an essential enzyme for synthetizing ceramide-based lipids, like CerPs^68^. Pharmacological modulation of LPS-treated BV-2 microglia with exogenous C2 ceramide demonstrated anti-inflammatory potential^69^, although its role in the endocytic pathway remains to be explored. Further, SM enrichment in M2a cells, coupled with more LEs in ARG1^+^ cells, is consistent with their role in endosomal maturation through neutral sphingomyelinase 2 (nSMase2)^70^, which facilitates the budding of intraluminal vesicles (ILVs) within late endosomes^70^.

Transitioning to pro-inflammation, sphingolipid metabolism is also state-specific, with almost no SMs found in M1 microglia. The observed SM depletion in the M1 state aligns with a previous study, supporting that LPS stimulation drives sphingosine-1-phosphate (S1P) synthesis at the expense of SM biosynthesis^71^. Additionally, a LysoPC/LysoPE-associated lipid shows M1-specific enrichment, in line with LPS-treated human and mouse macrophage lipidomes^56,72^, potentially underlying the enhanced cation signaling and cell body remodeling under pro-inflammatory conditions^73^. Overall, M1 microglia contained enlarged and denser lipid droplets and fewer late endosomes, a profile consistent with sustained pro-inflammatory signaling and inefficient lipid turnover^74,75^. While we had limitations in detecting neutral lipids, mainly stored in the LD core, the outer monolayer of LDs consists mainly of phospholipids, such phosphatidylinositols (PIs) and phosphatidylcholines (PCs)^76^. These species, particularly PCs, act as surfactants for LD stabilization^77^, further supporting the overall stable 3D architecture of LDs in pro-inflammatory microglia over time. Some phosphatidylcholines (PCs) have also been found relatively abundant in M1 cells, being consistent with their respective roles in pro-inflammatory signaling^78^.

Understanding the complex inter-organelle molecular communications and the high relevance of lipid species in promoting different cell states, our established 3D architecture approach targets a more global decoding of microglia states. This approach is inspired by how other omics analyses decipher cell populations in different organisms and outline intrinsic cell-to-cell variability within heterogeneous, unsorted populations^8^. The need for further time-course analyses to capture dynamic transitions calls for systems that fully recapitulate the diversity of primary microglia^79^. Through dimensionality reduction, these morphological units could potentially reveal structural heterogeneity across microglia, uncovering variability in polarization outcomes that extend beyond the classical M1/M2 spectrum^26^. Towards this direction, complementary nucleus 3D architecture analysis, including evaluation of heterochromatin-to-euchromatin composition and spatial distribution^24^, could also connect ultrastructure with epigenetic regulation, underpinning individual cell populations. Such work would also link cytoplasmic remodeling with transcriptional control mechanisms, such as histone modifications, already central to efforts promoting anti-inflammation in microglia^80,81^.

Collectively, our findings demonstrate that microglial polarization is accompanied by coordinated remodeling of ultrastructure and lipid composition. LD morphology and lipidomic profiles serve as sensitive markers of functional state^53,56^, providing mechanistic insights into the metabolic basis of microglial heterogeneity. These insights have translational relevance; targeting lipid species intervening in lipid droplet breakdown and endosomal maturation would increase precision in therapeutic interventions beyond systemic anti-inflammatory approaches. The overlap between M1 and aging-associated intracellular remodeling in microglia further highlights the potential for modulating metabolic and structural pathways in neurodegeneration^52^. Finally, this framework could be extended to other immune-relevant brain cell types, such as astrocytes and T cells, advancing our global understanding of intracellular instrumentation of ongoing neuroinflammation in neurodegenerative diseases.

## Resource availability

### Lead contact

Requests for further information and resources should be directed to and will be fulfilled by the lead contact, Venera Weinhardt.

### Materials availability

The BV-2 cell line, all materials and reagents used in this article are available upon request to the lead contact. Detailed protocols for experiments reported in this article are available upon request to the lead contact.

### Data and code availability

Original and analyzed images can be provided upon request to the lead contact. All codes related to 3D morphometric and correlated SXT-lipid analysis are reposited to GitHub and be accessed upon request. Any additional information needed to reanalyze the data is available upon request to the lead contact.

## Materials and Methods

### Cell culture & *in vitro* assays

The adult mouse microglia (BV-2) were purchased from Biological Resource Center (ICLC) cell bank (http://www.iclc.it/deposito.html; ICLC ATL03001). The cells were cultured continuously at 37°C with a 5% CO₂ supply in Dulbecco’s Modified Eagle Medium-High Glucose (9007.1, Roth), 10% Fetal Bovine Serum (FBS; F6765, Sigma Aldrich) and 1% Penicillin-Streptomycin (P4333, Sigma Aldrich). Cell density was maintained at 60-70% through subculturing twice a week, using 0.25% trypsin-EDTA (25200072, ThermoFischer Scientific). The BV-2 cells were polarized to M1 and M2a states by exposing them to 0.1-10 μg/mL Lipopolysaccharides (LPS) from Escherichia coli O127:B8 (L4516-1MG, Sigma Aldrich) or 20-40 ng/mL recombinant mouse interleukin-4 (IL-4; 404-ML-010, Biotechne) at varying concentrations in the growth medium for 6 or 24 h. Recovery from treatment was assessed by replacing the LPS- or IL-4- containing medium with a fresh growth medium and incubating the cells for an additional 24 h period. For M1 to M2a polarization, BV-2 cells were first challenged with LPS (10 μg/mL) for 24 hours, then incubated with fresh growth medium containing 10-200 ng/mL recombinant human sestrin 2 (SESN2; SRP0412-50UG, Sigma Aldrich) for another 24 hours.

### Western Blot

For Western blot analysis, 2 million cells from each condition were collected using trypsin-EDTA, washed with ice-cold PBS (1×), and lysed in 1% Triton X-100 with EDTA-free protease inhibitor cocktail (11836170001, Merck) in cold PBS. Lysates were centrifuged at 12,000 × g for 15 minutes at 4 °C, and supernatants were collected. Protein concentrations were measured using the Pierce™ BCA Protein Assay Kit (Thermo Fisher). Equal amounts of protein (15–20 μg per sample) were mixed with 2.5× denaturation buffer (5% SDS, 25% glycerol, 157.5 mM Tris-HCl, 0.08% bromophenol blue, β-mercaptoethanol), then boiled at 95 °C for 10 minutes. Proteins were separated on 7.5% SDS-PAGE gels (for iNOS, 130 kDa) or 10% gels (for ARG1, 35 kDa), and transferred to methanol-activated PVDF membranes (Millipore). Membranes were blocked with 5% BSA in TBS-T (0.1% Tween-20) for 1 hour at room temperature and incubated overnight at 4 °C with primary antibodies: rabbit anti-iNOS (1:1000; ab15323, Abcam) or rabbit anti-Arg1 (1:1000; ab91279, Abcam), diluted in blocking buffer. After washing, the membranes were incubated with mouse anti-α-tubulin (1:500; T9026, Sigma-Aldrich) for 1 hour at room temperature. Following further washes, the membranes were incubated for 1 hour with HRP-conjugated secondary antibodies, anti-rabbit and anti-mouse (1:10,000 each; 115-035-044, Jackson ImmunoResearch). Protein bands were visualized using the Pierce™ ECL Western Blotting Substrate (Thermo Fisher) and detected with an Advanced Fluorescence and ECL Imager.

### Organelle staining

Before *in vitro* polarization, BV-2 cells were seeded at 70–80% confluency either onto 12 mm round glass coverslips (fitted into 24-well plates) for confocal microscopy (protein detection) or into glass-bottomed 24-well plates for correlative wide-field microscopy. To visualize late endosomes (LEs), cells were incubated overnight at 37 °C in 5% CO₂ with CellLight™ Late Endosomes-GFP (30 particles per cell; C10588, Invitrogen) in their growth medium. For plasma membrane staining, cells were incubated with MemBrite® Fix 543/560 (1:1000 dilution; #30094-T, Biozol) for 5 minutes at 37 °C in 5% CO₂. Following live staining, cells were washed with pre-warmed 1× Hanks’ Balanced Salt Solution (HBSS) and fixed with 4% paraformaldehyde (PFA) for 15 minutes at room temperature, protected from light. After fixation, cells were washed with 1× PBS and incubated with DAPI (1:500; for nuclear staining, 6335.1, Roth) and HCS LipidTOX™ Red Neutral Lipid Stain (1:1000; H34476, Invitrogen) in 1× PBS for 1 hour at room temperature, protected from light. Following several PBS washes, stained cells on coverslips or in glass-bottomed plates were imaged using light microscopy. Notably, neither LipidTOX nor CellLight Late Endosomes-GFP signals were retained after permeabilization, making these markers well-suited for correlative analysis with immunofluorescence on the same cells (Figure SI4).

### Immunofluorescence

Following fixation with 4% PFA (and optionally organelle staining), immunostaining for iNOS or ARG1 was performed on cells seeded on coverslips or glass-bottomed plates. Cells were washed with PBS, permeabilized with 0.1% Triton X-100 in PBS for 5 minutes at room temperature and blocked with 1% BSA and 10% normal donkey serum in PBS-T for 1 hour. Primary antibodies (rabbit anti-iNOS, 1:100; or rabbit anti-Arg1, 1:200) were applied overnight at 4 °C in blocking buffer. The next day, the cells were washed and incubated with fluorophore-conjugated secondary antibodies (donkey anti-rabbit Alexa Fluor 647, 1:200, A31573, Invitrogen) and DAPI (1:500). After washing with PBS, the coverslips were mounted with antifade medium and sealed. Alternatively, the well plates were filled with PBS and sealed with parafilm. Samples were stored at 4 °C in the dark until imaging. Imaging was performed using a Leica TCS SP8 inverted confocal microscope with a 20×/0.75 oil immersion objective. Z-stack images were acquired using laser lines at 405, 532, and 638 nm. Detector settings were optimized using Leica software to avoid overexposure.

The confocal z-stack images were processed using Fiji software^82^. Maximum-intensity projections of each fluorescence channel were generated from the acquired z-stacks, with brightness and contrast adjusted for visualization purposes and background subtracted, using a 50-pixel radius. Organelle and immunofluorescence signals were automatically segmented for quantification using a custom Fiji Macro analysis script. Results of the segmentation are shown in Figure SI2. For single-cell quantification of iNOS⁺ and ARG1⁺ cells, only intact cells, identified by the presence of both a nucleus and an enclosing plasma membrane, were included; damaged or incomplete cells were excluded. Positive protein signals were defined as fluorescence localized specifically within the cytoplasmic area of intact cells. Manual quantification of these cells was performed using the multi-point tool in Fiji, with coordinates saved in the Region of Interest (ROI) Manager.

### Correlative fluorescence microscopy

Co-localization of organelles and proteins within the same cell population was assessed using a wide-field ACQUIFER imaging machine equipped with a 20× objective in brightfield and three fluorescence channels: DAPI (405 nm), GFP (488 nm), TRITC (550 nm), and Cy5 (633 nm). Initially, cells stained with LipidTOX™ Red (TRITC), CellLight™ Late Endosomes-GFP (GFP), and DAPI were imaged. To maintain consistent fields of view (FOVs), each well plate was precisely positioned on the sample stage, and the center of each well was determined based on the plate geometry. These coordinates were input into the ACQUIFER software to generate a unified image acquisition script. Multiple FOVs were acquired per well using an 8×8 grid centered on each well. Z-stacks were captured for each FOV using autofocus guided by the TRITC (LipidTOX) signal. Following the first imaging session, cells were processed for immunofluorescence staining of iNOS or ARG1. In the second imaging session, the TRITC and GFP channels were disabled due to the loss of LipidTOX and CellLight signals following permeabilization. Autofocus was instead based on the Cy5 channel for detection of iNOS or ARG1. The original acquisition script was reused to ensure identical FOVs and imaging settings across both sessions.

The images (.tiff) were organized as stacks, and then as hyperstacks, with each channel including the z-stacks of each field of view (FOV). Nuclei were used as fiducial markers for aligning and overlaying the FOVs (Figure SI4). The intact cells with overlayed nuclei, found in both imaging sessions, were chosen for the analysis. The selected cells containing organelles inside their cytoplasm were counted as LD+ or LE+. The iNOS+ or ARG1+ cells containing LDs or LEs were finally compared under different conditions. Quantification was described as described in the immunofluorescence analysis paragraph.

### Cryo-soft X-ray tomography

BV-2 cells, under control and LPS/IL-4-treated conditions, were harvested from culture after a short incubation with trypsin-EDTA, pelleted, and washed twice with 1x PBS. As a single-cell suspension, they were injected into glass capillaries and rapidly frozen by immersion in liquid propane cooled with liquid nitrogen (∼ −90°C)^83^. SXT data were acquired through full-rotation imaging using the soft X-ray microscope, XM-2, at the National Center for X-ray Tomography, located at the Advanced Light Source of Lawrence Berkeley National Laboratory in Berkeley, CA (https://ncxt.org/)^83^. Cells were exposed to a stream of liquid-nitrogen-cooled helium gas during the data collection process to prevent radiation damage^83^. Each dataset involved the collection of 184° rotation tomographs, with one projection image captured per 2° angle^83^. The automatic reconstruction software was used to reconstruct projection images into a 3D volume^83^. This process combined information from 92 projection images around the capillary at 184 degrees, followed by segmentation and 3D rendering, to visualize microglia in 3D^83^. LACs of individual lipid species were calculated, based on β values measured in Henke online calculator tool^57^ (density: -1 gm/cm^3^, photon energy: 520 eV and respective chemical formulas) and on the mathematical formula µ = 4rrf3/A, where μ is X-ray absorption coefficient (LAC) and λ attenuation length (nm)^84^.

The SXT datasets were processed with Amira 2020.3.1 software. First, the pixel intensities were corrected by multiplying them by the LAC value, known from the pixel size of each tomography dataset. The nucleus was segmented manually in every 10 virtual slices throughout the dataset, which was then interpolated to reconstruct the 3D labelled volume. The 3D labels from automatically segmented whole cells^41^, obtained using online Biomedisa tool^85^, were subtracted from nucleus 3D labels for semi-automatic threshold segmentation of mitochondria, lipid droplets, and endosomes, based on their individual LAC ranges.

### MALDI lipid mass spectrometry

For sample preparation, 3 cell pellets per condition were frozen and stored at −80 °C until measurement. Pellets were resuspended in 50 µL water by pipetting up and down three times. Three droplets (10 µL each) per treatment condition were then pipetted onto slides for measurement. A 1,5-diaminonaphthalene (DAN) matrix solution (10 mg mL⁻¹) was prepared in ACN/H₂O (60:40, v/v), sonicated for 15 minutes, and applied to the slides in eight spraying cycles using an M5 Sprayer (HTX Technologies LLC, Chapel Hill, USA). Spraying parameters were: 60% ACN in water, nozzle temperature 70 °C, bed temperature 35 °C, flow rate 0.07 mL min⁻¹, nozzle velocity 1200 mm min⁻¹, 2 mm track spacing in an HH pattern, spraying pressure 10 psi, gas flow rate 2 L min⁻¹, and a nozzle height of 40 mm above the slide surface. No dry time was included. After matrix application, slides were immediately transferred to the mass spectrometer. MALDI MS data was acquired using a timsTOF fleX from Bruker Daltonics (Bremen, Germany), with measurement settings being described in Table 2.

**Table 2.** MALDI lipid mass spectrometry settings using timsTOF fleX from Bruker Daltonics.

| Parameter | Positive Mode | Negative Mode |
| --- | --- | --- |
| Polarity | negative | positive |
| Range [$m/z$] | 300-2300 | 300-2300 |
| Laser shots | 400 | 400 |
| Laser frequency [1/s] | 10000 | 10000 |
| Laser field size [$\mu\text{m}$] | 50x50 | 50x50 |
| Funnel 1 RF [Vpp] | 420 | 420 |
| Funnel 2 RF [Vpp] | 400 | 400 |
| Multipole RF [Vpp] | 380 | 380 |
| Collision Energy [eV] | 10 | 10 |
| Collision RF [Vpp] | 1800 | 1800 |
| Low $m/z$ | 300 | 300 |
| Transfer Time [$\mu\text{s}$] | 105 | 85 |
| Pre Pulse Storage [$\mu\text{s}$] | 10 | 10 |

The timsTOF fleX MS imaging .tdf files were converted into .imzML files using timsconvert with 32-bit precision^86^, excluding ion mobility and without compression. The resulting .imzML files were uploaded to the false discovery rate (FDR) controlled METASPACE database for annotation using SwissLipids as the database^87^. Annotations with FDR ≤ 10% for both positive and negative ion modes were exported as .csv files to store all annotations. Putative sum formulas were used for lipid characterization. An in-house R script reduced the extracted .imzML files to the annotated peaks with a tolerance of ± 0.005 Da using the Cardinal R package for data storage prior to further processing^88^. These peaks were then compared for each treatment to the untreated control group, to identify significant differences using a one-sided t-test. Only the features with a significance over 0.05 were kept. The corresponding Cohen’s d effect sizes were calculated for each group using the same script. Results from positive and negative ion mode were merged for the final plot and sorted according to their corresponding LAC Absorption.

### MALDI trapped ion mobility spectrometry (TIMS) MS Imaging and lipid identification by prm-PASEF

Molecules annotated on MS1 level via METASPACE database with FDR ≤ 10% were analyzed on MS2 level using parallel reaction monitoring-parallel accumulation serial fragmentation (prm-PASEF) Scan mode as follows: First, MALDI trapped ion mobility spectrometry (TIMS) MSI runs were set up on MS1 level to extract molecule-specific mobility windows [V*s/cm^2^] for mobility-based isolation. MALDI TIMS-MSI was conducted on a timsTOF flex system (Bruker Daltonics) equipped with a smartbeam 3D 10kHz laser and TimsControl 6.1 and flexImaging v7.6 software (Bruker Daltonics). Data was acquired in positive and negative ion mode with the settings described in Table 3, and the TIMS settings detailed in Table 4.

**Table 3.** MALDI trapped ion mobility spectrometry (TIMS) MSI settings for positive and negative ion mode data acquisition.

| Parameter | Positive Mode | Negative Mode |
| --- | --- | --- |
| Polarity | positive | negative |
| Range [$m/z$] | 300-1800 | 300-1800 |
| Laser shots | 240 | 240 |
| Laser frequency [1/s] | 5000 | 5000 |
| Laser field size [$\mu\text{m}$] | 50x50 | 50x50 |
| Funnel 1 RF [Vpp] | 500 | 500 |
| Funnel 2 RF [Vpp] | 500 | 500 |
| Multipole RF [Vpp] | 600 | 600 |
| Collision Energy [eV] | 10 | 10 |
| Collision RF [Vpp] | 1800 | 1800 |
| Low $m/z$ | 300 | 300 |
| Transfer Time [$\mu\text{s}$] | 80 | 80 |
| Pre Pulse Storage [$\mu\text{s}$] | 5 | 5 |

**Table 4.**
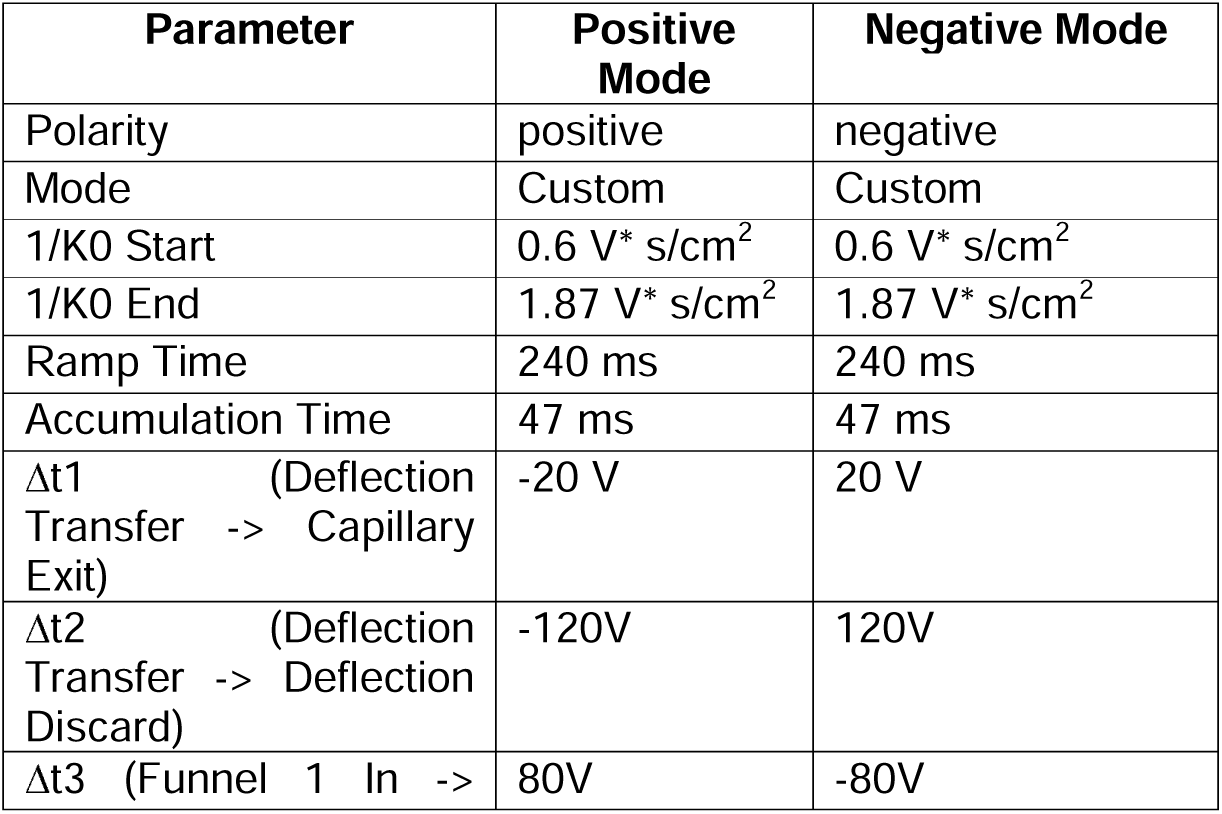
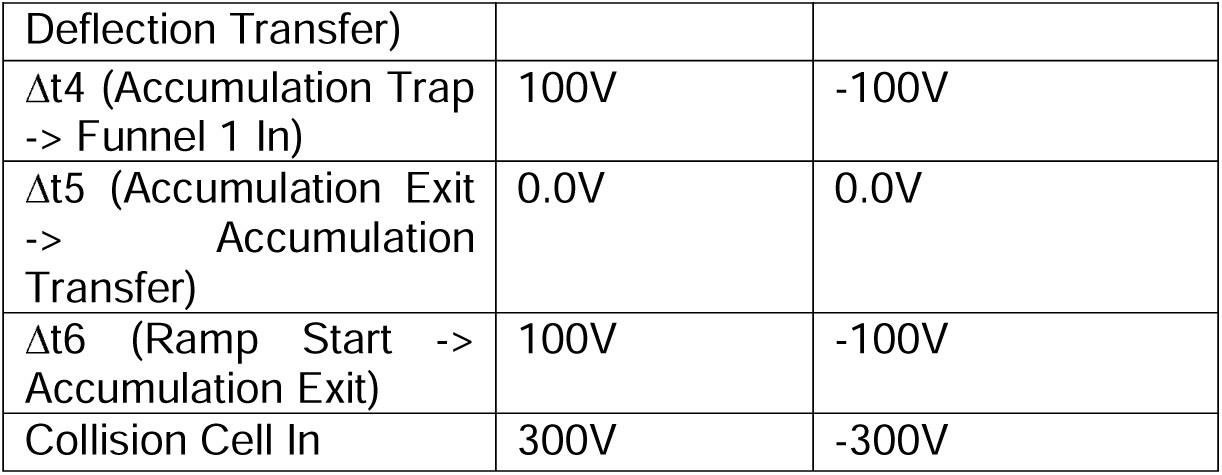
Additional TIMS-MSI settings for positive and negative ion mode data acquisition.

External mass and ion mobility calibration was done via ESI source, using ESI-Low Concentration Tuning Mix (Agilent Technologies, Santa Clara, USA). Mass calibration was done in Enhanced Quadratic mode and mobility calibration was done in Linear mode. .mis files were uploaded in SCILS software v.5.1 (Bruker Daltonics) to extract molecule specific mobility information (1/K0 V*s/cm^2^) stored in .spp files for the following prm-PASEF analysis. prm-PASEF analysis was done using mobility-based isolation and fragmentation settings. Mobility-based separation information was stored in .spp files. Each molecule was separately fragmented due to overlapping mobility windows. Several parameters were adjusted for screening low mass fragment ions in negative and positive ion mode, as described in Table 5.

**Table 5.**
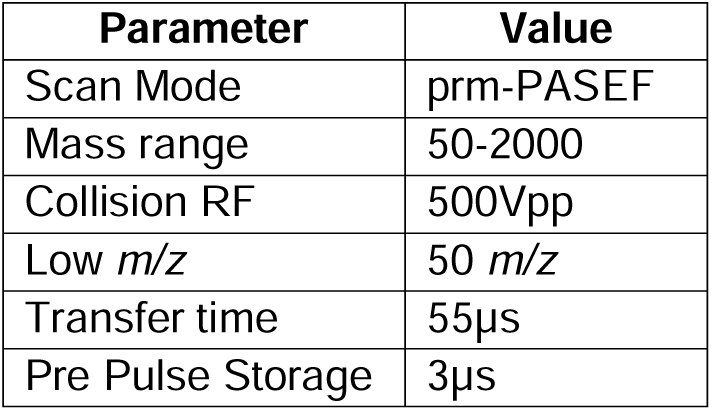
Low mass fragment ion screening settings in negative and positive ion mode using prm-PASEF analysis.

| Parameter | Value |
| --- | --- |
| Scan Mode | prm-PASEF |
| Mass range | 50-2000 |
| Collision RF | 500Vpp |
| Low $m/z$ | 50 $m/z$ |
| Transfer time | 55μs |
| Pre Pulse Storage | 3μs |

Isolation width settings were set as follows: for *m/z* 400 a width of 2 *m/z* and for *m/z* 1100 a width of 4 *m/z*. Collision Energy settings were experimentally adjusted between 30-90 eV using 0.1 mobility steps 1/K0 [V*s/cm^2^] as base. Exact Collision Energies [eV] used for each molecule can be found in Figures SI6-13. Spectra were acquired by accumulating 30 shots on bulk cells.

### High mass MALDI trapped ion mobility spectrometry (TIMS) MSI with prm-PASEF

TIMS measurements of molecules with a mass ≥ 1400 *m/z* were conducted with reduced gas flow (2.1 mbar) using a mass range from 300-2500 m/z and mobility range of 1-2.2 V*s/cm^2^. Collision RF was adjusted to 2000 Vpp, Transfer Time was set to 110µs and Pre Pulse Storage to 10 µs. For the prm-PASEF analysis of molecules with a mass ≥ 1400 *m/z* following parameters were adjusted: mass range from 200-2500, Collsion RF 1500 Vpp, Transfer Time 90µs, Pre Pulse Storage 8 µs.

### Spectra analysis

prm-PASEF spectra were analysed using DataAnalysis software v6.2 (Bruker Daltonics). MassWiki (MassWiki) and Lipid Maps (LIPID MAPS) were used for spectra comparison and annotation. Chemical structures were drawn using the open source software ChemSketch (Free Chemical Drawing Software for Students | ChemSketch | ACD/Labs).

### Statistical analysis

Data analysis and graphical data visualization were performed in Python with Jupyter Notebook^89^, utilizing data frames imported from Excel files (.csv). The sample size (n), t-test, and Pearson’s correlation analysis are provided in figures and figure legends, accordingly.

## Acknowledgments

V.W. was funded for the current work within the framework of the Excellence Strategy of the Federal and State Governments of Germany, the Excellence Cluster “3D Matter Made to Order” (3DMM2O), the CLEXM MSCA-DN project funded by the European Union under Horizon Europe, and by the CoCID project (no. 101017116) funded within the EU Research and Innovation Act. A.C. conducted the experiments at NCXT, with support from the Joachim Herz Stiftung Foundation’s program for ‘Add-on Fellowships for Interdisciplinary Life Science’, cohort 2022, and Boehringer Ingelheim Fonds. C.L. and the NCXT were supported by NIH NIGMS P30GM138441 and DOE Biological and Environmental Research Project DE-AC02-05CH11231. C.H. was supported by SFB1638 project “lipid imaging and lipidomics” with number 511488496. Prof. Joachim Wittbrodt’s laboratory and the BioQuant building at Heidelberg University provided laboratory space and technical support for the current work. NCXT at LBNL also provided technical and laboratory support to A.C. to conduct the SXT imaging experiments.

## Author Contributions

V.W. and A.C. conceived the presented idea of whole-cell architecture investigation in neuroinflammatory microglia. A.C. performed the molecular characterization assays for *in vitro* polarization. X.P. assisted with Western blot assays. A.C., supervised by J-H., and V.L., performed the SXT imaging. A.C. developed the presented correlative fluorescence pipeline and consequent light microscopy analysis. V.W. and A.C. conceived the 3D morphometric SXT analysis. A.E. assisted with 3D analysis by providing the ACSeg model. J.H. and J.L.C. performed the mass spectrometry characterization and analyzed the data. M.A.L.G., C.L., C.H., and V.W. supervised the presented work and provided research infrastructure. All authors equally contributed to the preparation of the final manuscript.

The core content of this manuscript is derived from A.C.’s PhD thesis, which was submitted and defended at Heidelberg University in February 2025. The thesis is available in the university’s library archive upon request, yet this manuscript includes revised and updated content.

## Supplemental Information

**Table SI1.**
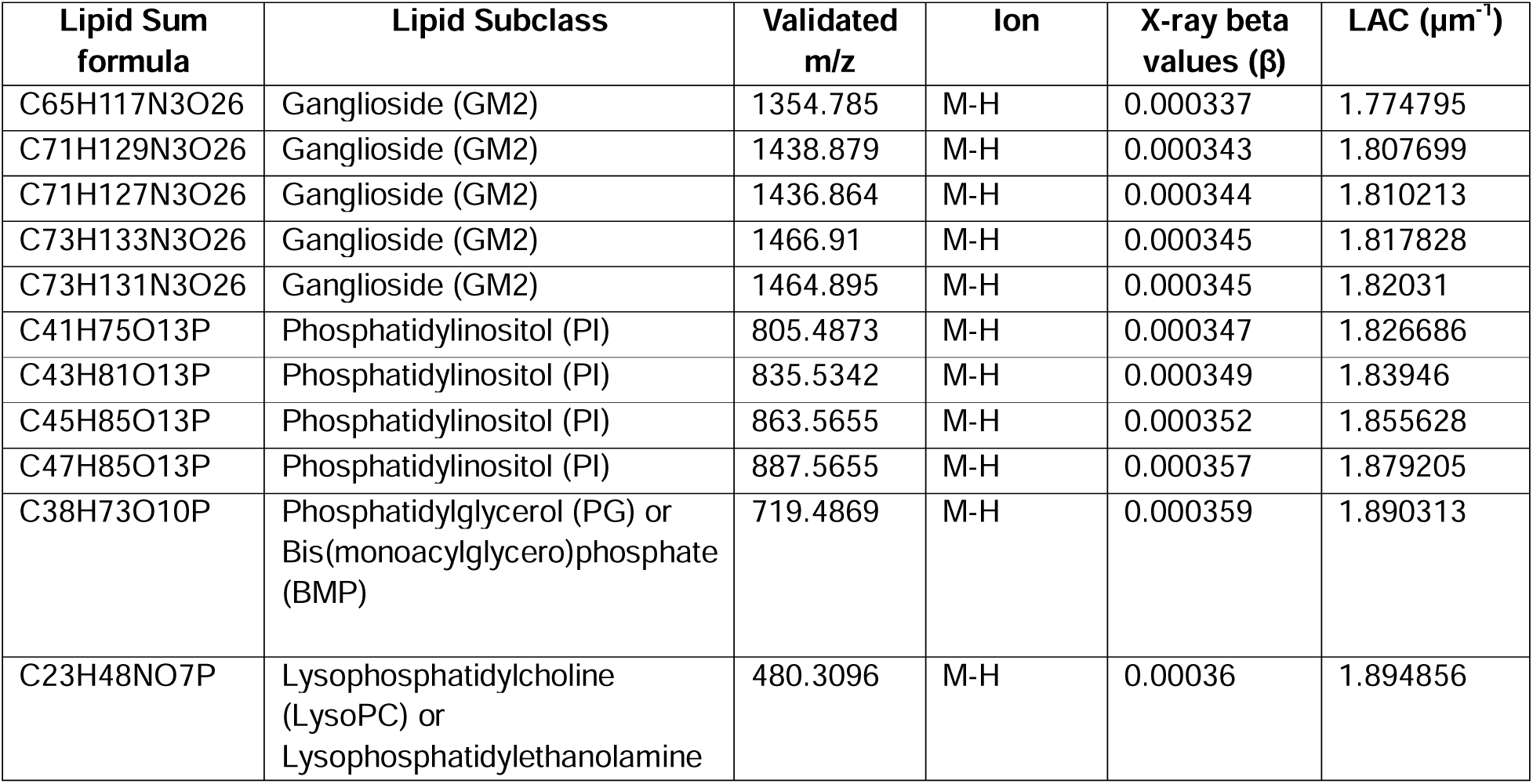
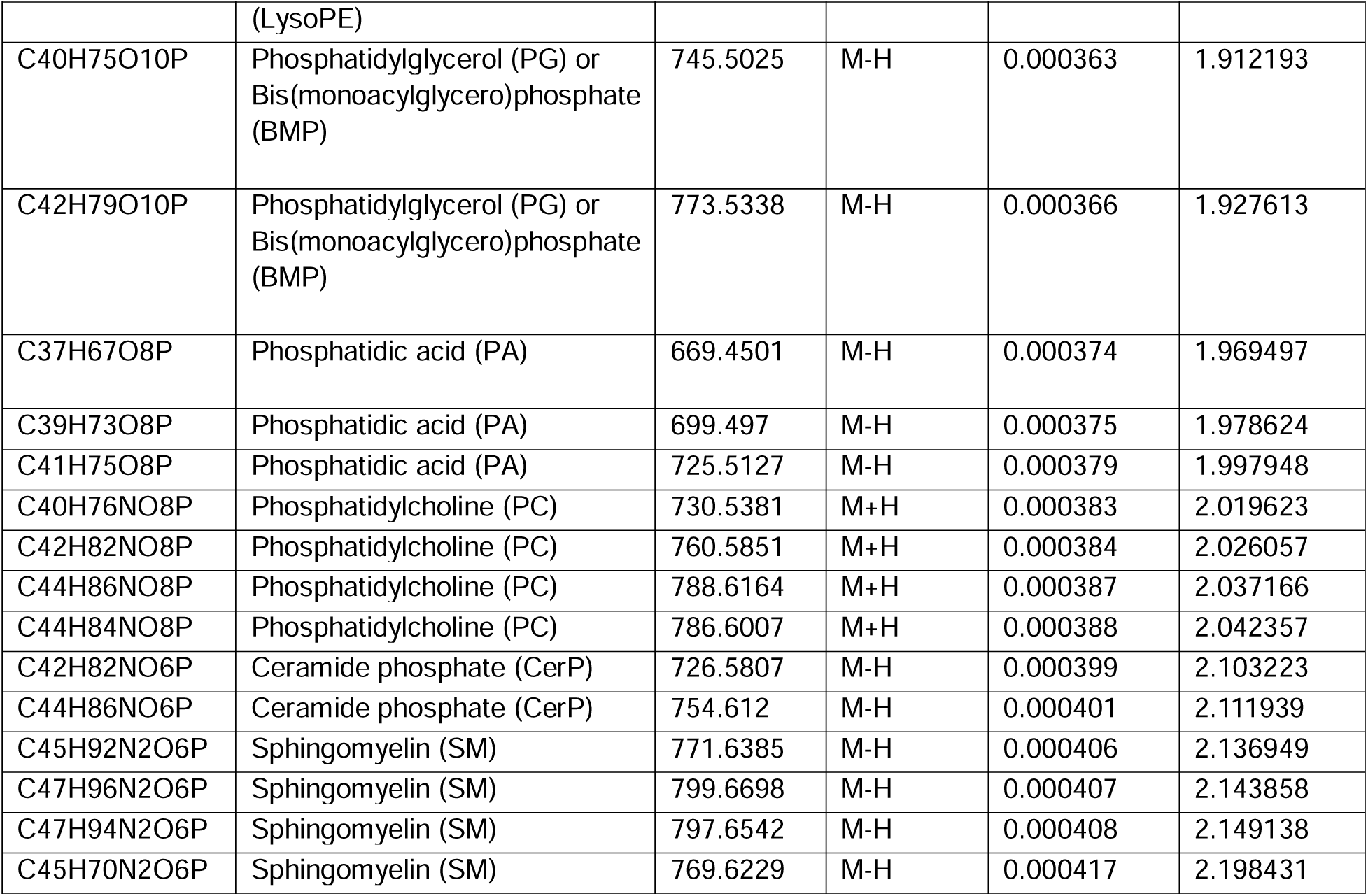
SXT-mass spectrometry analysis on 26 validated lipid species. *density is -2.2 g/cm^3^ by default from Henke calculator (https://henke.lbl.gov/optical_constants/getdb2.html)

**Figure SI1.**
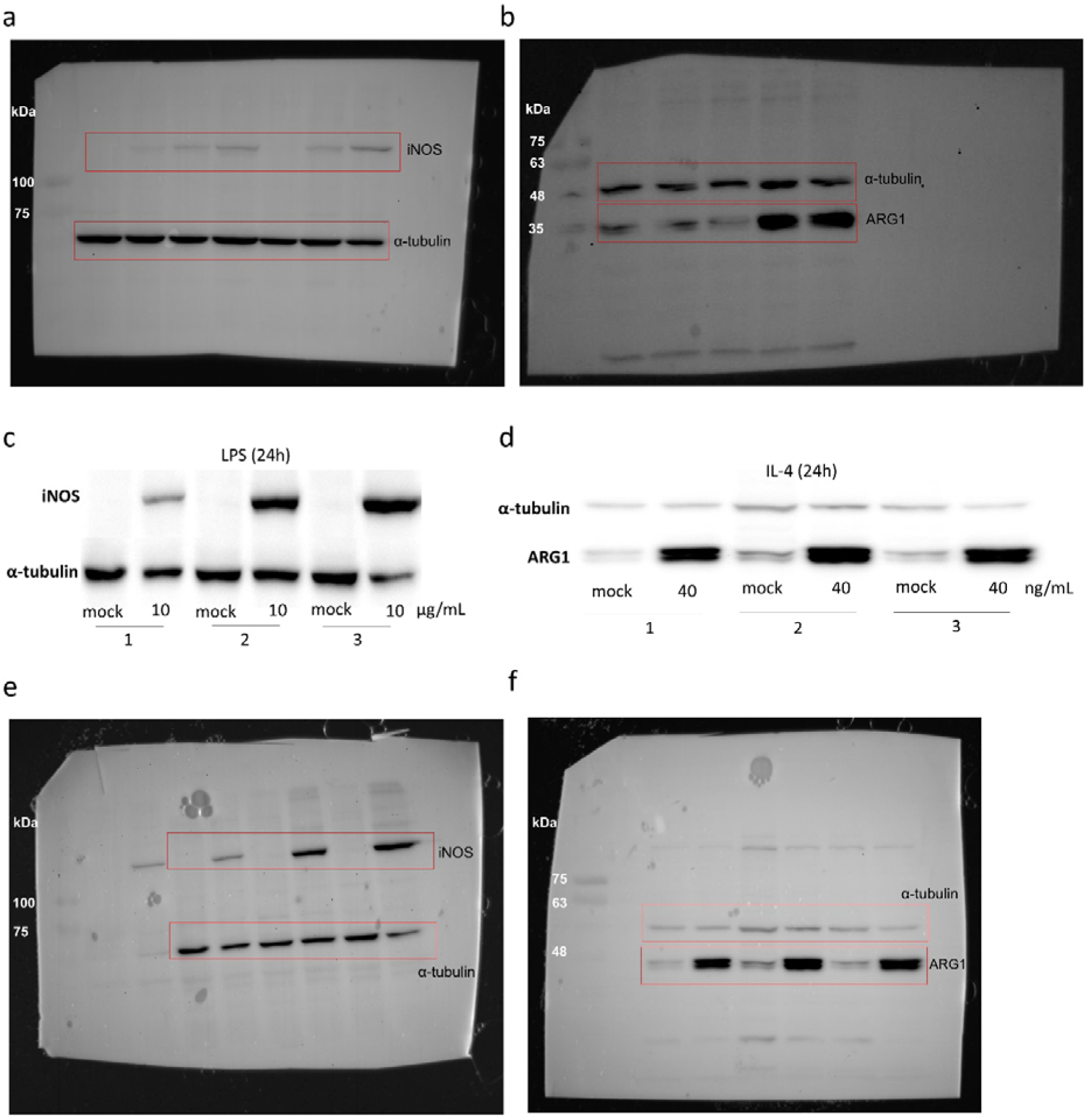
Western blot images related to Figure 1 and examination of *in vitro* BV-2 robustness **(a-b)** The whole images of the Western blot membranes for iNOS (130 kDa, 8% Tris-Glycine gel) and ARG1 (35 kDa, 12% Tris-Glycine gel) detection with the protein ladder in the first well. Red boxes refer to Figure1, panel b images. **(c-d)** Three independent *in vitro* activation experiments of BV-2 with LPS and IL-4 were measured for average iNOS and ARG1 expression relative to α-tubulin (55 kDa), respectively. **(e-f)** The whole images of Western blot membranes for the three independent activation assays from panels c and d, showed in red boxes, respectively.

**Figure SI2.**
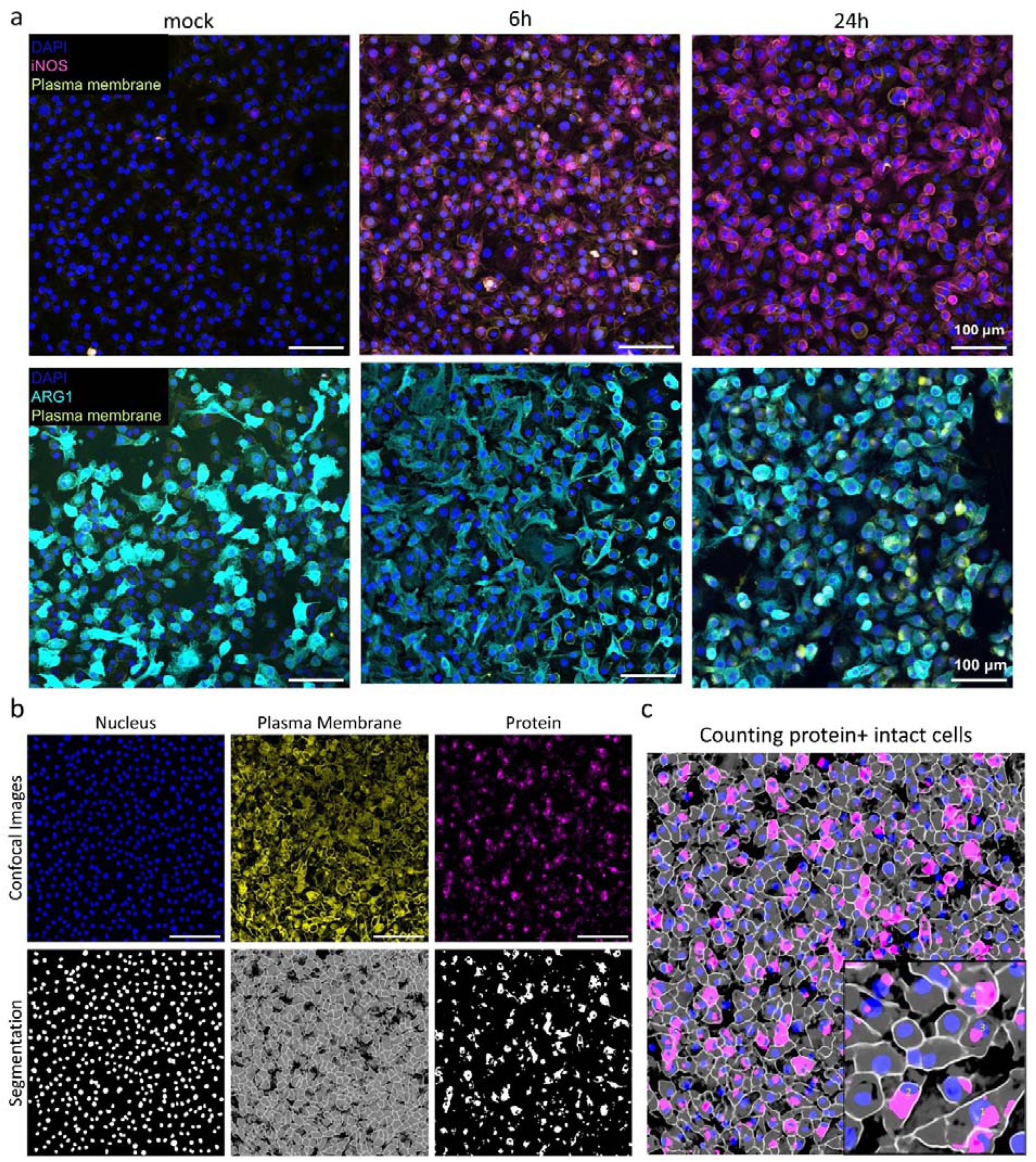
Immunofluorescence images and analysis related to Figure 1 **(a)** Z-stack projection of confocal image with all respective channels (nucleus-blue, plasma membrane-yellow, and immunofluorescent protein-magenta (iNOS) or cyan (ARG1)). The scale bar is 100 μm. **(b)** Z-stack projection of confocal image with all respective channels (nucleus, plasma membrane, and immunofluorescent protein) with their segmentation results. The scale bar is 150 μm. **(c)** Quantification process based on intact cell counting.

**Figure SI3.**
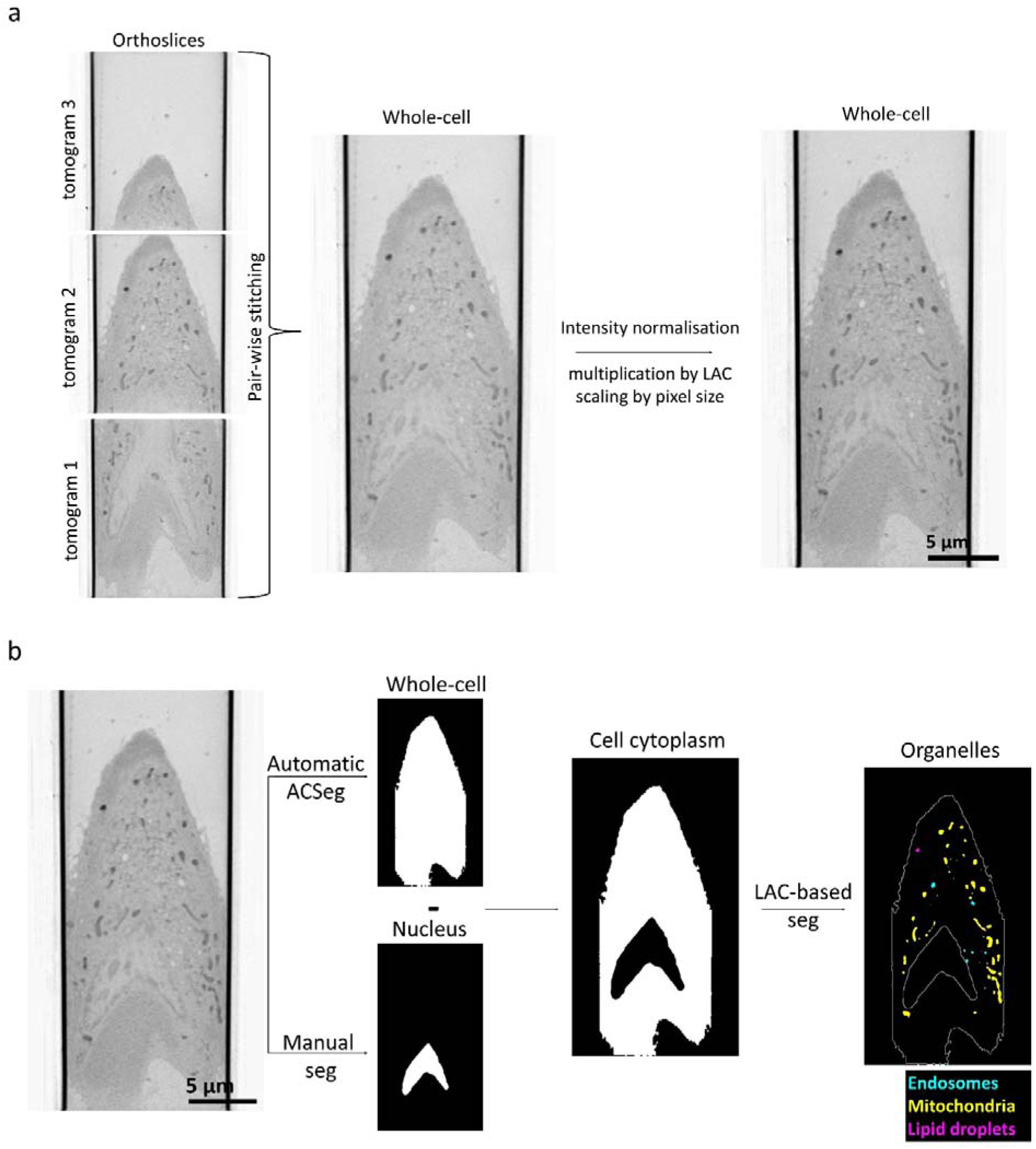
SXT image processing and segmentation related to Figure 2 **(a)** Tomogram stitching using a pair-wise approach to view a whole cell within one FOV. The stitched tomogram is then normalized for its pixel intensity and size by the LAC factor and pixel size, respectively. **(b)** Threshold-based segmentation of individual organelles lying in the cytoplasm volume requires the subtraction of the automatically segmented whole-cell from the manually segmented nucleus, to remove interfering LAC values of the nucleus volume and background noise.

**Figure SI4.**
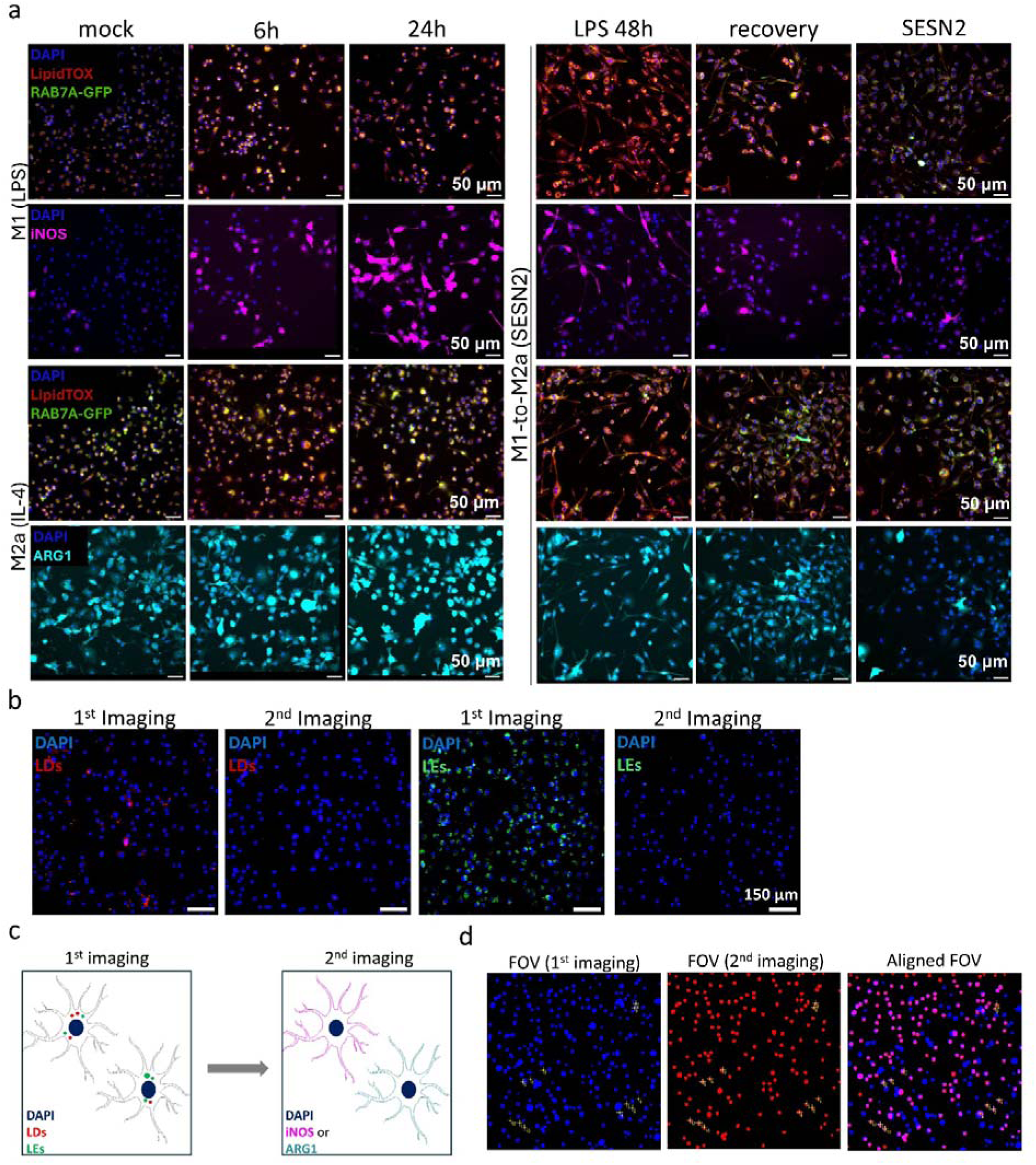
Correlative fluorescence imaging for organelle and protein markers related to Figure 3 **(a)** Sequentially acquired images showing organelles (lipid droplets – LipidTOX in red, late endosomes – Rab7A-GFP in green, nucleus – DAPI in blue) localized in activated (iNOS – magenta, ARG1 – cyan) BV-2 cells under M1, M2a and M1-to-M2a transition. Immunofluorescent images from second imaging have been transferred in x and y to show overlapping areas. The scale bar is 50 μm. **(b)** Sequential wide-field imaging on mock BV-2 cells before and after immunofluorescence. The organelle markers for LDs (red) and LEs (green) are faded because of permeabilization. **(c)** Schematics show the correlative fluorescence imaging workflow. **(d)** Nuclei (DAPI) are the fiducials markers for overlapping the organelles to proteins of each cell imaged in first (blue) and second (red) field of view (FOV). Due to further processing of cells with immunostaining, some of the cells are washed out as seen of the aligned FOV (magenta shows overlap).

**Figure SI5.**
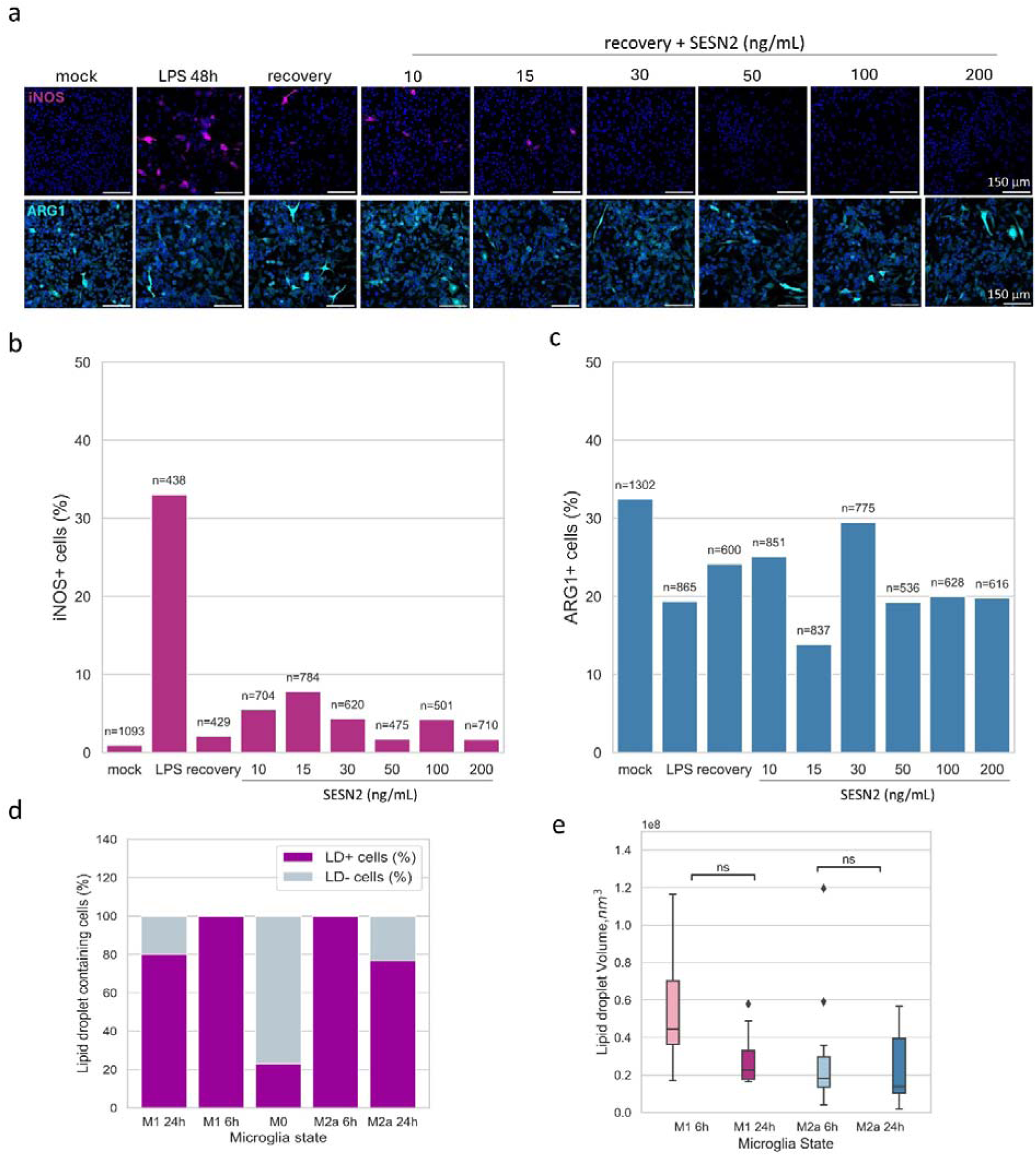
M1-M2a promotion related to Figure 3 and 3D lipid droplet analysis related to Figure 4 **(a)** Confocal images were acquired with an oil-immersed 20x objective, and the scale bar is 150 μm. Single-cell protein expression is quantified, based on their cytoplasmic intensity, for **(b)** iNOS^+^ and ARG1^+^ cell proportions. Quantification was based on two confocal images per condition. The sample size (n=cells) per group for iNOS^+^ quantification is mock (no LPS) n=1093, 48h (LPS) n=438, 48h (24h recovery) n=429, 48h (24h SESN2 10 ng/mL) n=704, 48h (24h SESN2 15 ng/mL) n=784, 48h (24h SESN2 30 ng/mL) n=620, 48h (24h SESN2 50 ng/mL) n=475, 48h (24h SESN2 100 ng/mL) n=501 and 48h (24h SESN2 200 ng/mL) n=710. The sample size (n=cells) per group for ARG1^+^ quantification is mock (no LPS) n=1302, 48h (LPS) n=865, 48h (24h recovery) n=600, 48h (24h SESN2 10 ng/mL) n=851, 48h (24h SESN2 15 ng/mL) n=837, 48h (24h SESN2 30 ng/mL) n=775, 48h (24h SESN2 50 ng/mL) n=536, 48h (24h SESN2 100 ng/mL) n=628 and 48h (24h SESN2 200 ng/mL) n=616. **(d)** M1 and M2a share an equal slope in lipid droplet (LD)-containing dynamics, compared to M0. **(e)** LD volume does not change significantly at M1 and M2a over time.

**Figure SI6.**
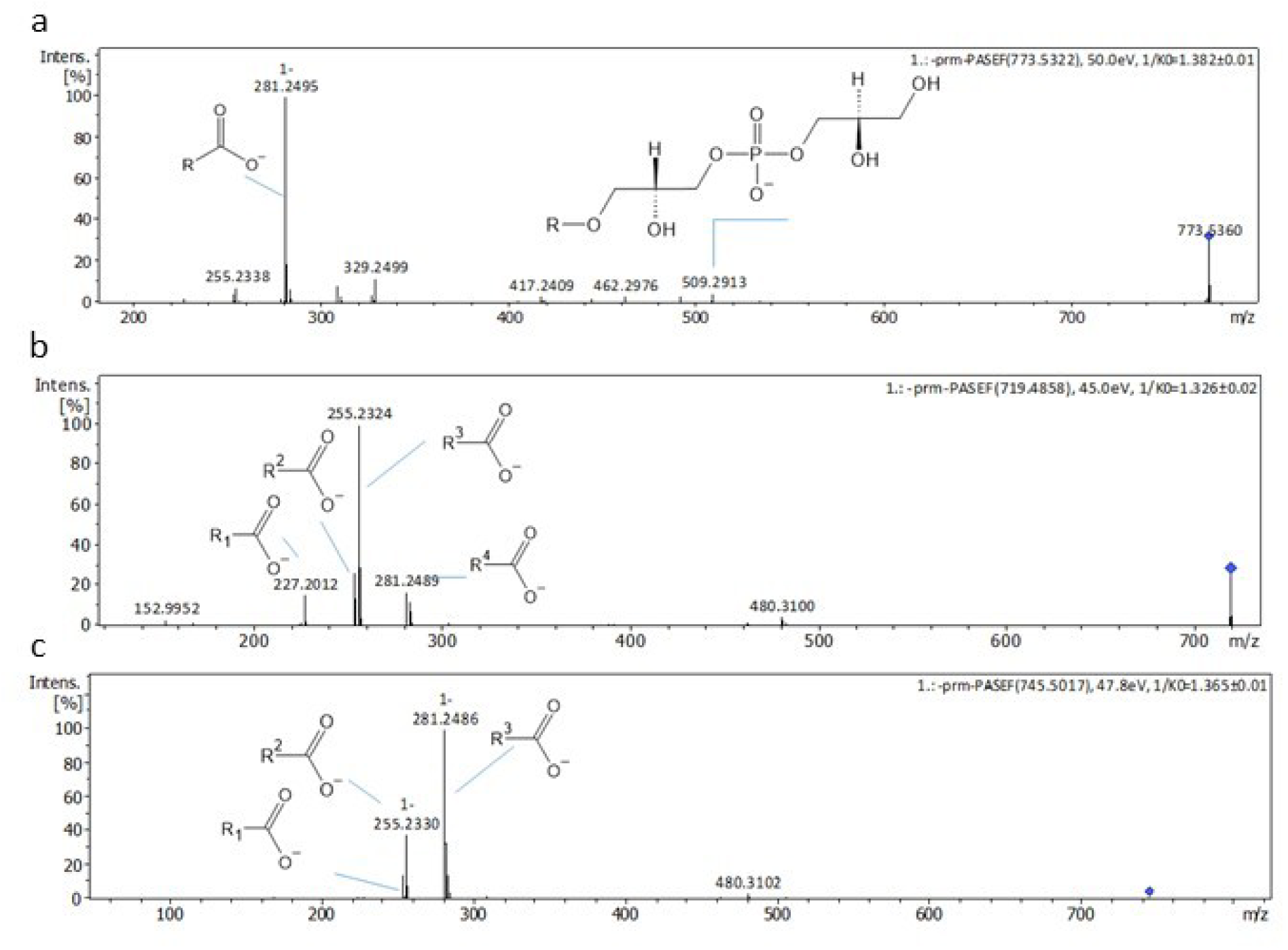
MS2 validation of PG/BMP related to Figure 4 In negative ion mode molecular ions [M-H]- were matched with the following sum formulas C42H79O10P (773.53 m/z), C38H73O10P (719.49 m/z) and C40H75O10P (745.50 m/z), on MS1 level this could be annotated to BMP/PG with an FDR of 0.05-0.1. PG and BMP are isobaric and cannot be distinguished on MS2 level in negative ion mode. **(a)** MS/MS spectra of molecular ion 773.53 m/z, BMP/PG 36:2, with R=FA18:1. **(b)** MS/MS spectra of molecular ion 719.49 m/z, BMP/PG 32:1 with R1 = FA14:0, R2 = FA16:1, R3 = FA16:0 and R4 = FA18:1. BMP/PG 14:0/18:1 and BMP/PG 16:0/16:1 have the same mass on MS1 level and could be both identified on MS2 level. **(c)** MS/MS spectra of molecular ion 745.50 m/z, BMP/PG 34:2 with R1 = FA16:1, R2 = FA16:0 and R3 = FA18:1. R1 and R3 indicate for the presence of BMP/PG 16:1/18:1, the presence of R2 let assume that BMP/PG 16:0/18:2 might be also present, although FA18:2 (279.23 m/z) could not be detected.

**Figure SI7.**
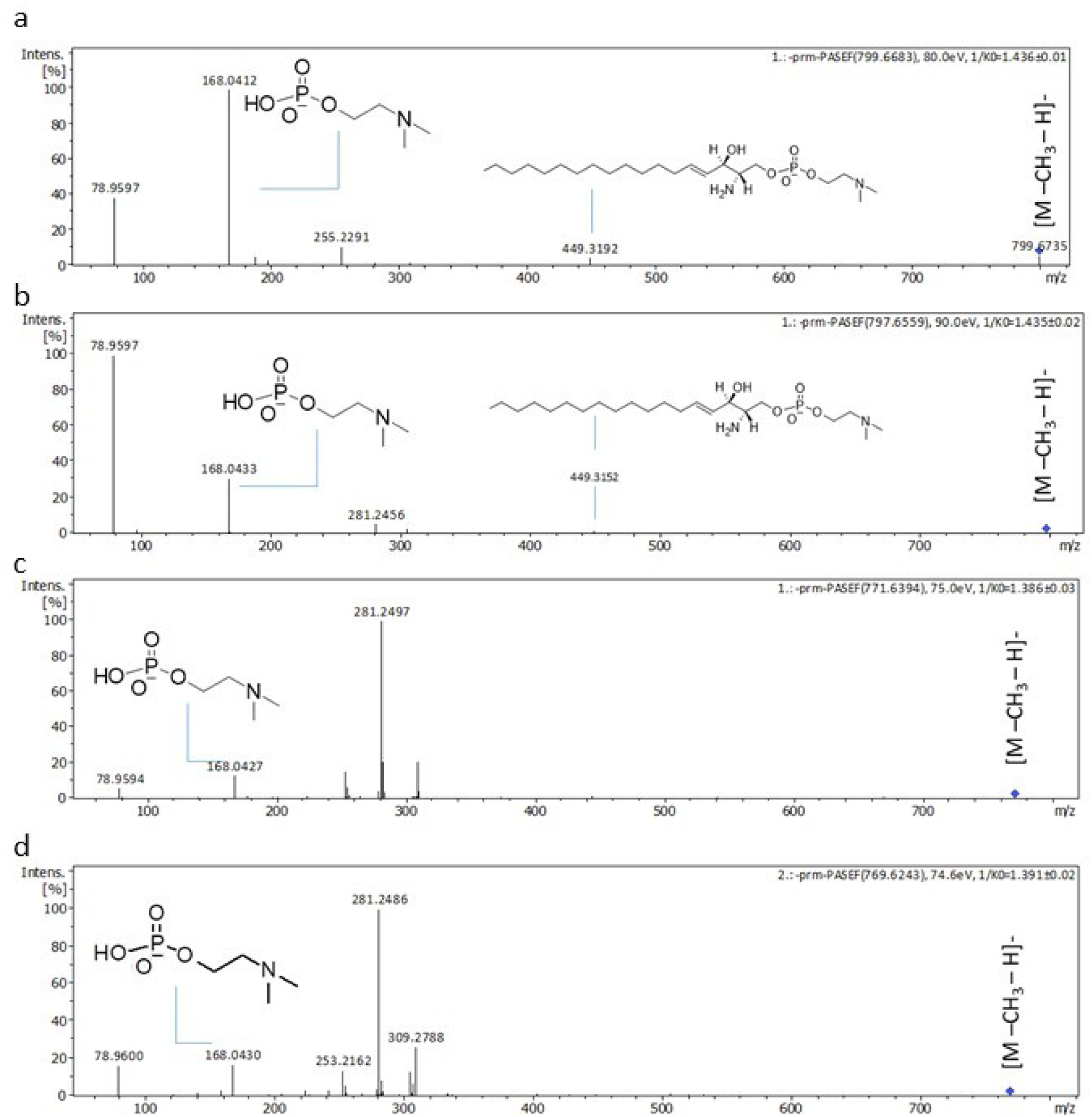
MS2 of SM related to Figure 4 In negative ion mode molecular ions [M-H]- 799.69 m/z, 797.66 m/z, 771.64 m/z and 769.62 m/z were assigned to be either CER-PE or SM on MS1 level with FDR 0.05-0.1. However CER-PE, a sphingolipid found mostly in invertebrates, is not commonly detected in mammalian cells^1^ and therefore unlikely. Furthermore, presence of CER-PE would be indicated on the MS2 level by the presence of the phosphoethanolamine head group fragment (140.07 m/z), which was not detected. In negative ion mode, SM are detected as [M-CH3-H]- molecular ions, a consequence of in-source demethylation^2^. Consequently, MS/MS spectra of molecular ions were screened for the demethylated phosphocholine head group fragment (168.04 m/z). **(a)** MS/MS spectra for molecular ion peak C47H96N2O6P (799.66 m/z), SM 42:1;O2. **(b)** MS/MS spectra for molecular ion peak C47H94N2O6P (797.66 m/z), SM 42:2;O2. **(c)** MS/MS spectra for molecular ion peak C45H92N2O6P (771.64 m/z), SM40:1;O2. **(d)** MS/MS spectra for molecular ion peak C45H70N2O6P (769.62 m/z), SM 40:2; O2.

**Figure SI8.**
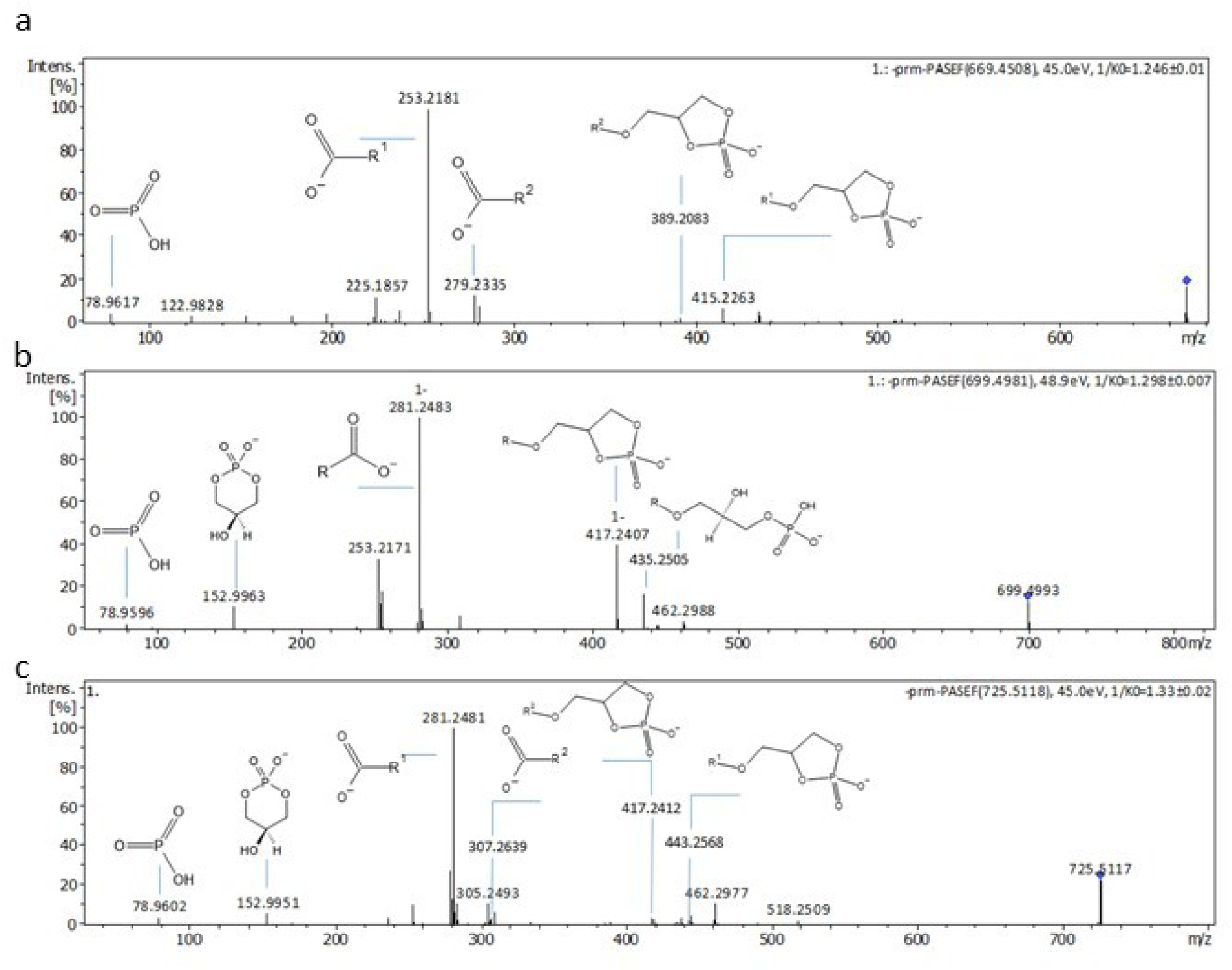
MS2 of PA related to Figure 4 In negative ion mode, molecular ions [M-H]- were matched with the following sum formulas C37H67O8P (669.45 m/z), C39H73O8P (699.50 m/z) and C41H75O8P (725.51 m/z). On MS1 level these sum formulas were annotated as PA with an FDR of 0.1. **(a)** MS/MS spectra for molecular ion peak 669.45 m/z [M-H]-, PA 34:3, with R1 = FA16:1 and R2 = FA18:2. **(b)** MS/MS spectra for molecular ion peak 699.49 m/z [M-H]-, PA 36:2, R = FA18:1. **(c)** MS/MS spectra for molecular ion peak 725.51 m/z [M-H]-, PA 38:3, R1 = FA18:1, R2 = FA20:2.

**Figure SI9.**
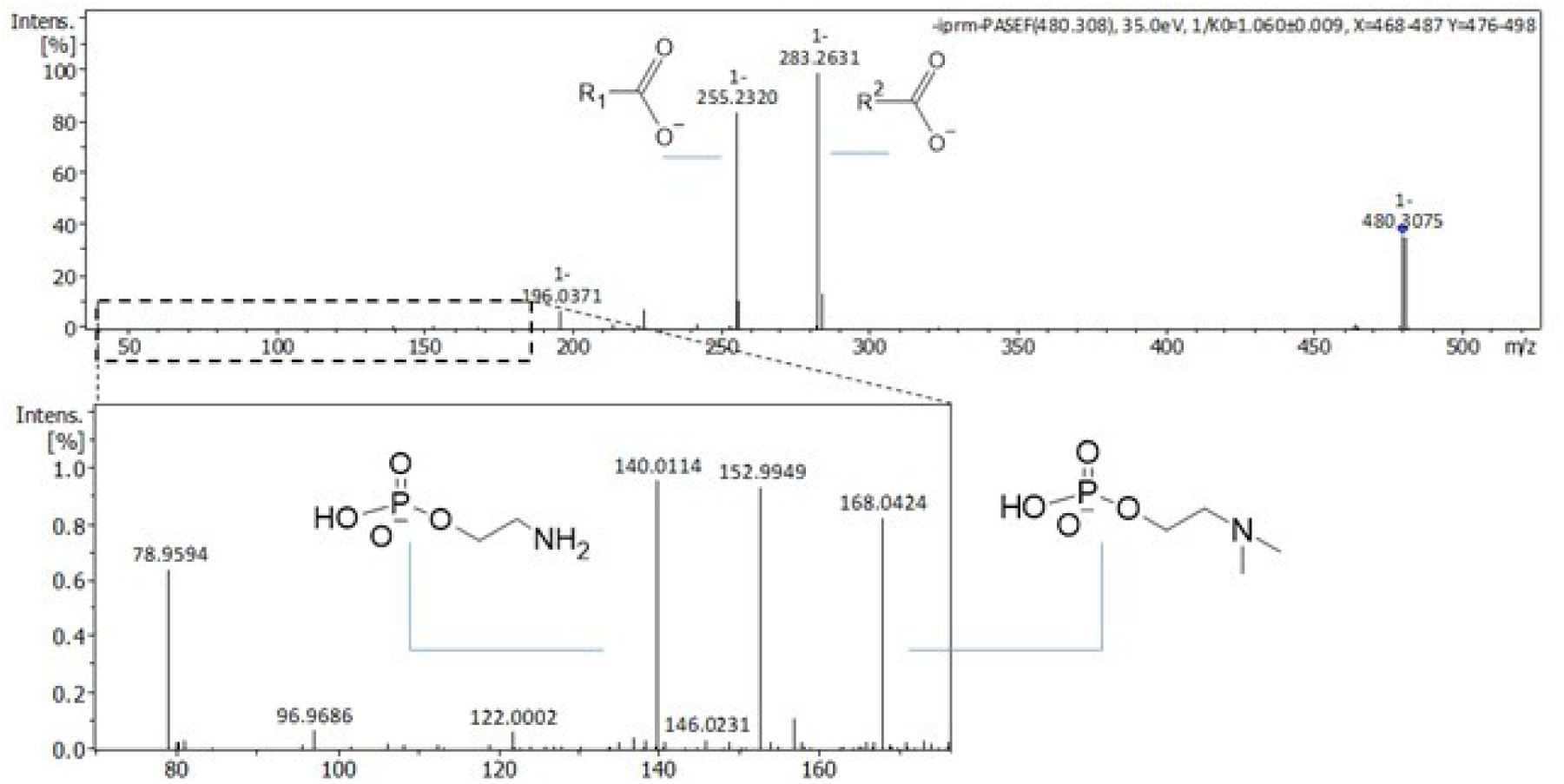
MS2 of LysoPE/LysoPC related to Figure 4 In negative ion mode, the molecular ion [M-H]- 480.31 m/z was matched to the sum formula C23H48NO7P with FDR = 0.1 on MS1 level. MS/MS analysis revealed a chimeric spectrum containing both LysoPC 16:0 (R1 = FA16:0) and LysoPE 18:0 (R2 = FA18:0). LysoPC is detected in negative ion mode as [M – CH3 – H]- molecular ion. Both headgroup fragments ethanolamine phosphate (140.01 m/z) and the phosphocholine group with CH3 loss (168.04 m/z) could be detected.

**Figure SI10.**
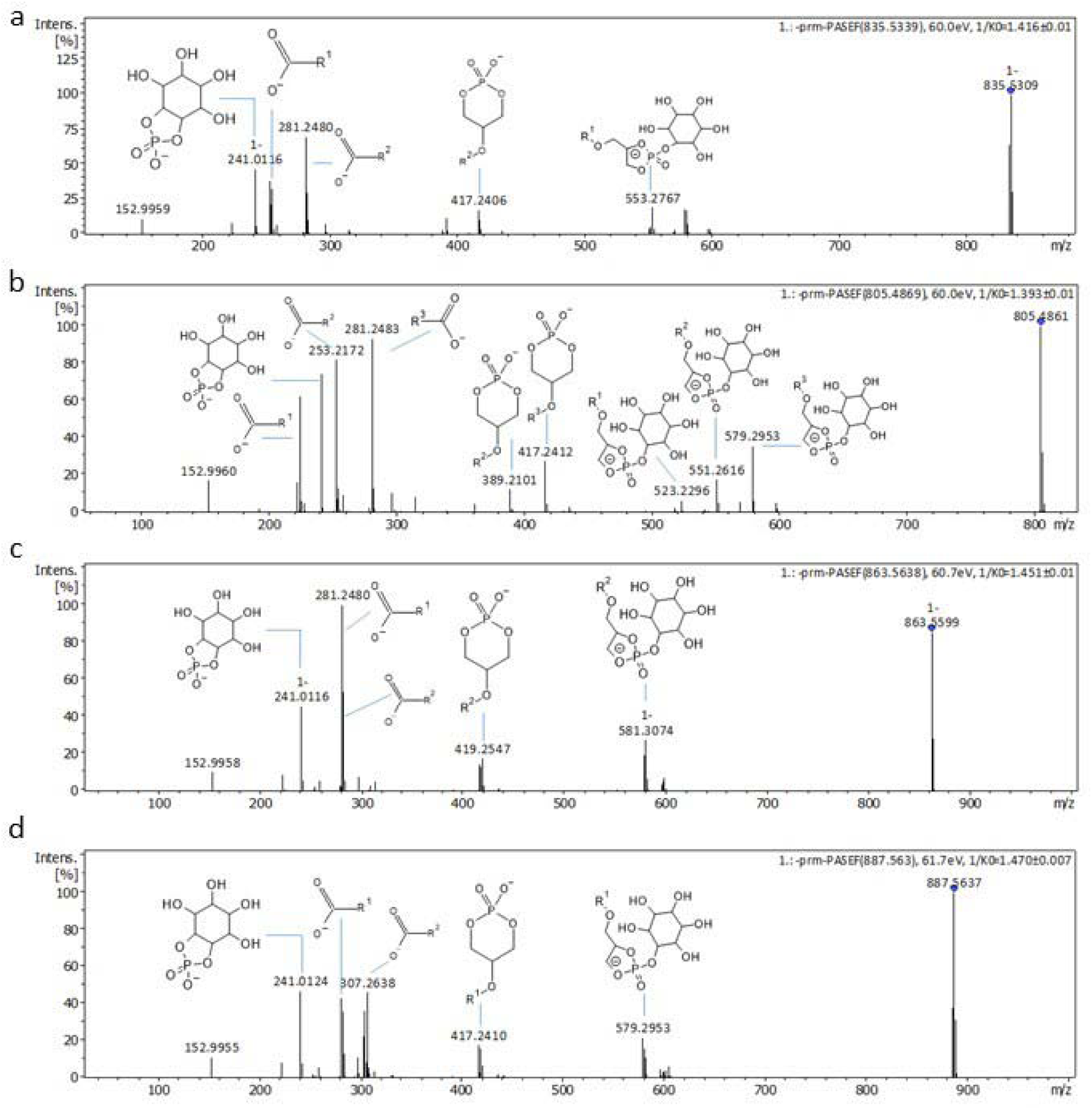
MS2 of PI related to Figure 4 In negative ion mode, molecular ions [M-H]^-^ were matched to the following sum formulas C43H81O13P (835.53 *m/z*), C41H75O13P (805.49 *m/z*), C45H85O13P (863.57 *m/z*), C47H85O13P (887.57 *m/z*) and identified as PI on MS1 level with FDR 0.05-0.1. MS2 spectra were screened for the inositol phosphate – H_2_O ion (241.01 *m/z*) as signature fragment for PI in negative ion mode. **(a)** MS/MS spectra for molecular ion [M-H]^-^ 835.53 *m/z*, PI 34:1, with R_1_ = FA16:1 and R_2_ = FA18:1. **(b)** MS/MS spectra for molecular ion [M-H]^-^ 805.49 m/z, PI 32:2, containing both PI 14:1/18:1 and PI16:1/16:1, with R_1_ = FA14:1, R_2_ = FA16:1 and R_3_ = FA18:1. **(c)** MS/MS spectra for molecular ion [M-H]^-^ 863.56 *m/z*, PI 36:1, with R_1_ = FA18:1 and R_2_ = FA18:0. **(d)** MS/MS spectra for molecular ion [M-H]^-^ 887.56 *m/z*, PI 38:3, with R_1_ = FA18:1 and R_2_ = FA20:2.

**Figure SI11.**
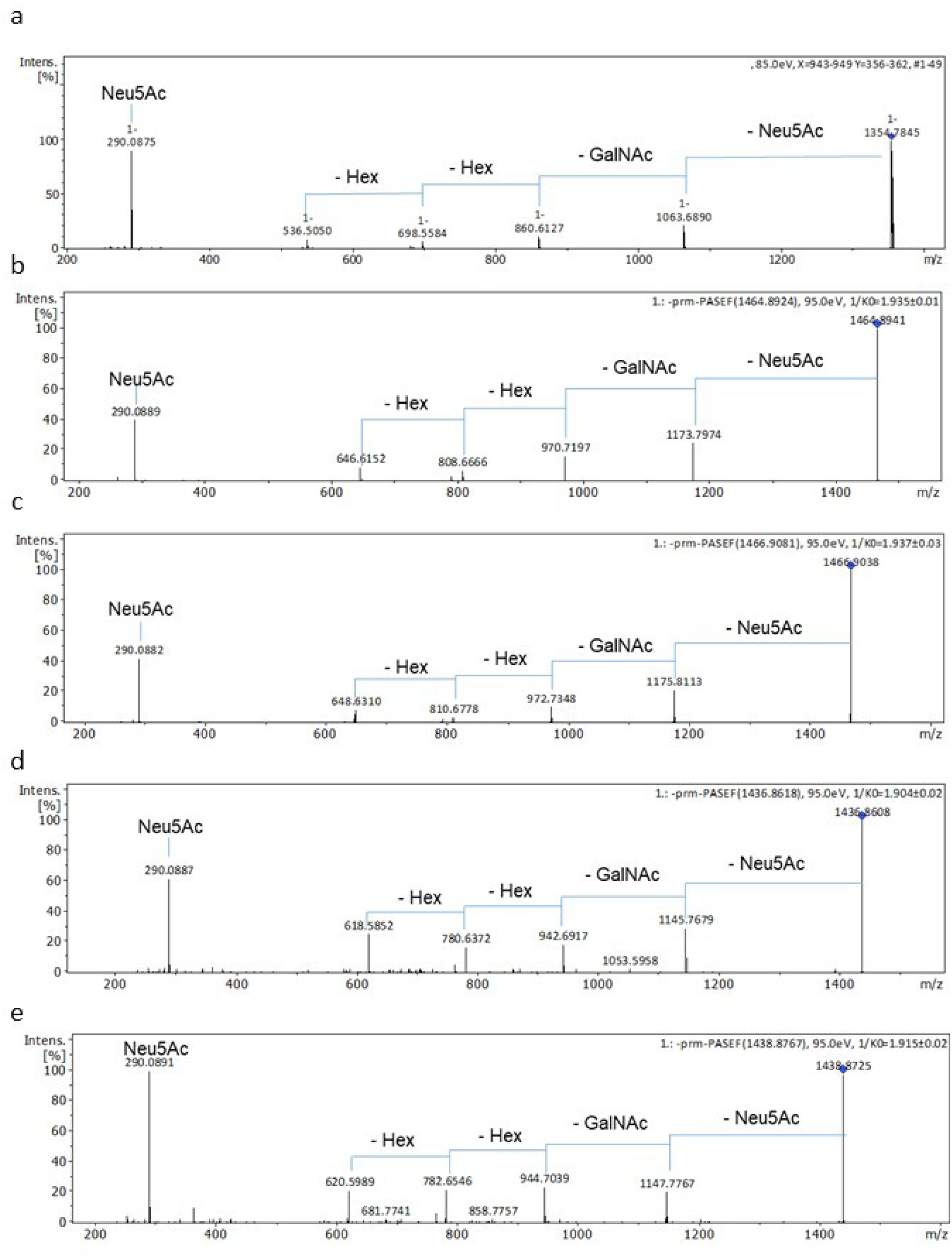
MS2 of GM related to Figure 4 In negative ion mode, molecular ions were matched to the following sum formulas C65H117N3O26 (1354.79 *m/z*), C73H131N3O26 (1464.89 *m/z*), C73H133N3O26 (1466.91 *m/z*), C71H127N3O26 (1436.86 *m/z*) and C71H129N3O26 (1438.88 *m/z*). On MS1 level these molecules were assigned to GM2 with a FDR 0.05-0.1. **(a)** MS/MS spectra of molecular ion 1354.79 *m/z,* GM2α (d34:1) **(b)** MS/MS spectra of molecular ion 1464.89 *m/z*, GM2α (d42:2). **(c)** MS/MS spectra of molecular ion 1466.91 *m/z*, GM2α (d42:1) **D** MS/MS spectra of molecular ion 1436.86 *m/z,* GM2α (d40:2). **(e)** MS/MS spectra of molecular ion 1438.88 *m/z,* GM2α (d40:1). Comparing MS/MS spectra to literature^3^ the characteristic fragmentation pattern could be identified with -291.1 Da (loss of Neu5Ac), -203.08 Da (loss of GalNAc) and -162.05 Da (loss of Hex).

**Figure SI12.**
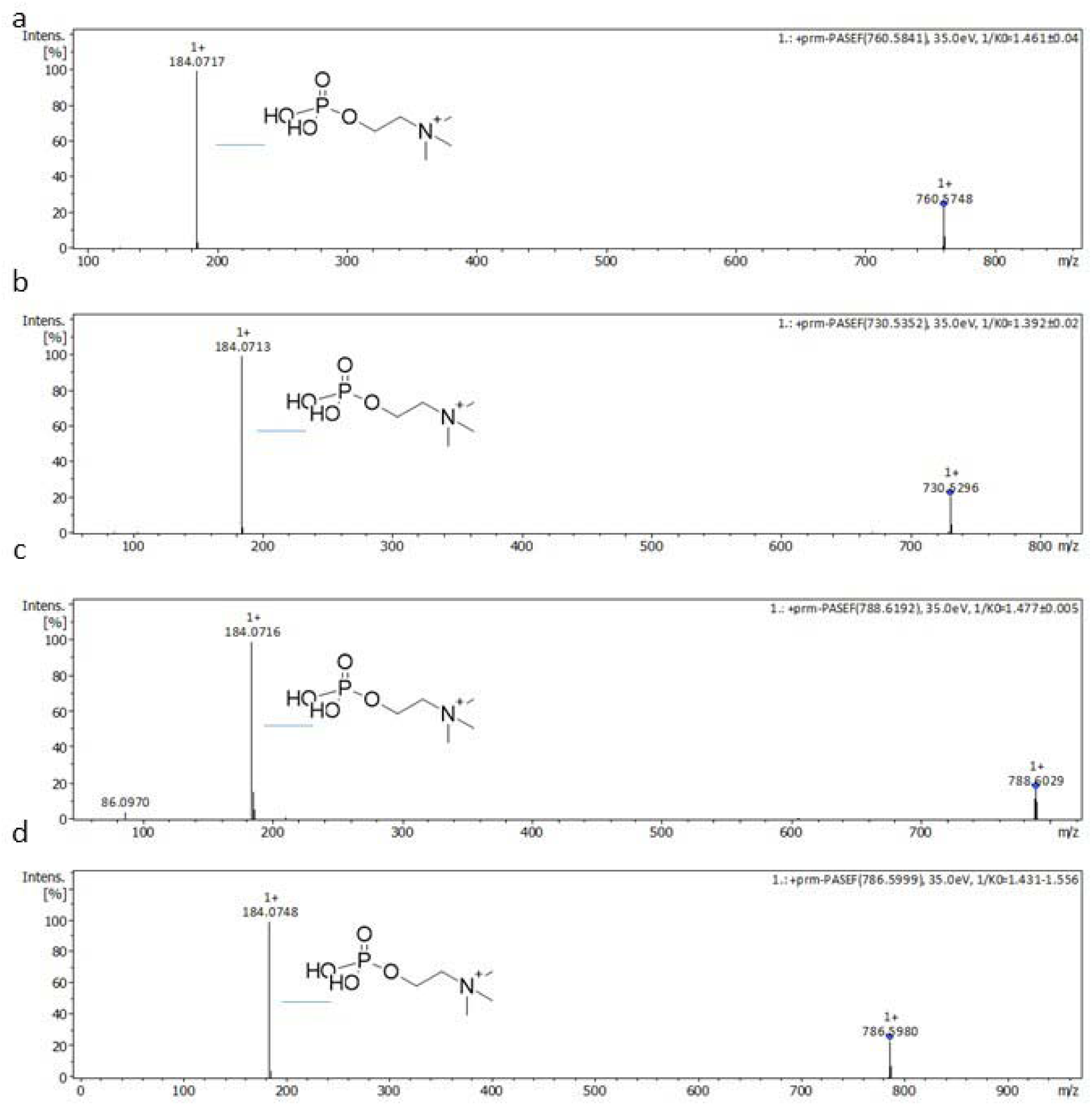
MS2 of PC related to Figure 4 In positive ion mode, molecular ions [M+H]^+^ were matched with the following sum formulas C42H82NO8P (760.58 *m/z*), C40H76NO8P (730.53 *m/z*), C44H86NO8P (788.6029 *m/z*) and C44H84NO8P (786.5980 *m/z*). On MS1 level these molecular ions were assigned as either PC or PE. **(a)/(b)/(c)/(d)** MS/MS spectra could identify for all 4 molecular ion peaks the phosphocholine head group (184.07 *m/z*) indicating for a PC lipid class. In positive ion mode PE would be detected with a neutral loss of the phosphoethanolamine head group (– 141.02 Da), which could not be detected here and thereby ruled out.

**Figure SI13.**
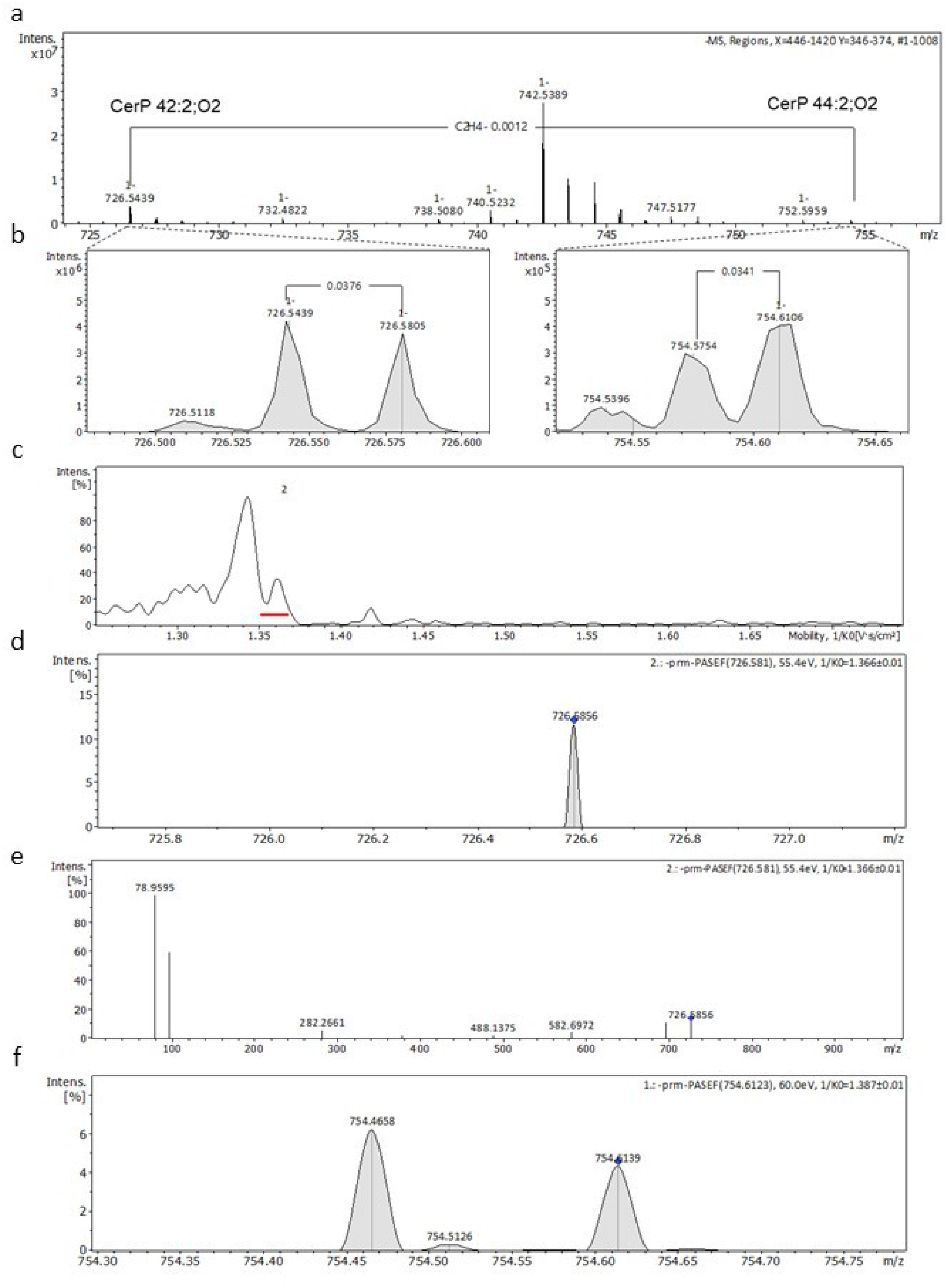
MS2 of CerP related to Figure 4 In negative ion mode, molecular ions [M-H]^-^ were matched to the following sum formulas C42H82NO6P (726.58 m/z) and C44H86NO6P (754.61 m/z). They were assigned to CerP on MS1 level with a FDR of 0.05-0.1. **(a)** On MS1 level 726.58 *m/z* and 754.61 *m/z* were identified as the molecular ion peaks [M-H]^-^ of CerP 42:2;O2 and CerP 44:2; O2, respectively. Based on MS1 level annotation, the assigned molecules differ in the presence of a C_2_H_4_ group. This could be detected on MS1 level, as indicated by a 28.03 Da mass shift. **(b)** Magnification revealed close proximity (Δ m/z = 0.0376) to an abundant molecular ion peak, thus making isolation of the molecular ion peaks 726.58 *m/z* and 754.61 *m/z* and thereby MS/MS analysis difficult. This observation was already described in the literature^4^. **(c)/(d)** Based on ion mobility the molecular ion peak 726.58 *m/z* could be isolated. **(e)** The corresponding MS/MS spectrum shows the most prominent fragment for CerP in negative ion mode ([M-H]^-^, 78.95 *m/z*)^5^. Comparing this to the literature, 726.58 m/z is the molecular ion [M-H]^-^ of CerP 18:1;O2/24:1^5^. **(f)** Ion mobility-based separation did not lead to an isolation for molecular ion peak 754.61 m/z, thereby MS/MS analysis was not possible. However, on MS1 level the molecular ion [M-H]^-^ 754.61 m/z could be assigned to CerP 18:1;O2/26:1, in comparison to literature^5^.

